# A multi-omics analysis of glioma chemoresistance using a hybrid microphysiological model of glioblastoma

**DOI:** 10.1101/2022.10.29.514383

**Authors:** Shahla Shojaei, João Basso, Meitham Amereh, Javad Alizadeh, Tania Dehesh, Simone De Silva Rosa, Courtney Clark, Misha Hassan, Mateuz Tomczyk, Laura Cole, Grant Hatch, Vern Dolinsky, Chris Pasco, David Schibli, Sanjiv Dhingra, Abhay Srivastava, Amir Ravandi, Rui Vitorino, Saeid Ghavami, Mohsen Akbari

**Affiliations:** Department of Mechanical Engineering, University of Victoria, Victoria, BC, Canada; Department of Human Anatomy and Cell Sciences, University of Manitoba, Winnipeg, MB, Canada; Faculty of Pharmacy, University of Coimbra, Coimbra, Portugal; Coimbra Chemistry Centre, Department of Chemistry, University of Coimbra, Coimbra, Portugal; Department of Biostatistics and Epidemiology, School of Public Health, Kerman University of Medical Sciences, Kerman, Iran; Department of Pharmacology and Therapeutics, University of Manitoba, Winnipeg, Canada; Department of Physiology and Pathophysiology, University of Manitoba, Winnipeg, Manitoba, Canada; University of Victoria-Genome BC Proteomics Centre, Victoria, BC, Canada; Physiology and Pathophysiology, University of Manitoba, St. Boniface Hospital Research Center, Winnipeg, Canada; iBiMED-Department of Medical Sciences, University of Aveiro, Aveiro, Portugal; UnIC, Department of Surgery and Physiology, Faculty of Medicine, University of Porto, Porto, Portugal; QOPNA & LAQV-REQUIMTE, Chemistry Department, University of Aveiro, Aveiro, Portugal; Research Institute of Oncology and Hematology, Cancer Care Manitoba, University of Manitoba, Winnipeg, MB, Canada; Faculty of Medicine in Zabrze, University of Technology in Katowice, Academia of Silesia, 41-800 Zabrze, Poland; Centre for Advanced Materials and Related Technologies, University of Victoria, Victoria, BC, Canada; Terasaki Institute for Biomedical Innovation, Los Angeles, United States; School of Biomedical Engineering, University of British Columbia, Vancouver, BC, Canada; Biotechnology Center, Silesian University of Technology, Akademicka 2A, 44-100, Gliwice, Poland

## Abstract

Chemoresistance is a major clinical challenge in the management of glioblastoma (GBM) Temozolomide (TMZ) is the chemotherapeutic drug of choice for GBM; however, the therapeutic effect of TMZ is limited due to the development of resistance. Recapitulating GBM chemoresistance in a controlled environment is thus essential in understanding the mechanism of chemoresistance. Herein, we present a hybrid microphysiological model of chemoresistant GBM-on-a-chip (HGoC) by directly co-culturing TMZ-resistant GBM spheroids with healthy neurons to mimic the microenvironment of both the tumor and the surrounding healthy tissue. We characterized the model with proteomics, lipidomics, and secretome assays. The results showed that our artificial model recapitulated the molecular signatures of recurrent GBM in humans. Both showed alterations in vesicular transport and cholesterol pathways, mitotic quiescence, and a switch in metabolism to oxidative phosphorylation associated with a transition from mesenchymal to amoeboid. This is the first report to unravel the interplay of all these molecular changes as a mechanism of chemoresistance in glioblastoma. Moreover, we have shown that the acquisition of resistance increases invasiveness and the presence of neurons decreases this property.

## Introduction

Glioblastoma (GBM) is the most aggressive and deadliest type of malignant central nervous system tumors with a median survival rate of 15 months [1]. The standard of care for GBM (the Stupp protocol) involves surgical resection of tumor to the extent possible, followed by radiotherapy and temozolomide (TMZ) chemotherapy. This approach has remained unchanged for the last decades and is ineffective, primarily due to the development of drug-resistant cell populations [2]. Chemoresistance in glioma cells is a complex process that involves multiple factors including dysregulation of programmed cell death pathways [3], presence of tumor residues that contain cancer stem cell (CSC) populations [4, 5], and the activation of DNA repair system. With high plasticity from a dormant status to highly proliferative, depending on the tumor microenvironment, [6, 7], activation of DNA repair systems [8–10], and modulation of autophagic and apoptotic cell death [11–13], CSC are referred to as the source of chemoresistance. However, the exact mechanism of resistance is not clear and understanding mechanisms underlying drug resistance and identifying novel modalities of therapy has remained as the major focus of research in the field [14].

Despite tremendous efforts in developing new therapeutics for GBM, most drugs have failed to improve the overall survival of patients in clinical trials. Poor clinical outcomes of developed drugs are primarily attributed to the inability of existing preclinical models to emulate the pathophysiology of GBM [15]. Current models for delineating glioma formation and progression rely on two-dimensional (2D) monolayer culture systems [16] or xenograft and genetically engineered models [17]. Two-dimensional (2D) cultures fail to recapitulate the complex GBM tumor microenvironment (TME), including the blood-brain-barrier and the extracellular matrix (ECM) that acts as a barrier to drug diffusion [18]. Apart from cost, anatomical and physiological differences with human, and ethical concerns for *in vivo* models [19], the use of nude rodents to develop xenograft models and histologically modified mice for genetic engineering models diverge tumor microphysiology from human glioblastoma. [20, 21].

Three-dimensional (3D) culture systems are alternatives to traditional 2D culture models as they enable recreating the complexities of tumor microenvironment more precisely. In this regard, multicellular tumor organoids are the simplest 3D culture systems that are produced by self-aggregation of cells in non-adherent wells to simulate cell-cell adhesion, diffusive gradients of nutrients and waste within solid tumors, and cell heterogeneity. The role of diffusive nutrient gradients in driving cancer stem cells with TMZ resistance that leads to the overexpression of DNA repair genes has been shown and the inner necrotic core of tumors has been introduced as the cancer stem cell niche [7, 22]. However, spheroids lack vasculature and interactions between tumors and the surrounding tissue [23]. To address above challenges, multi-compartment microfluidic systems have been used where tumor and non-tumor cells are cultured in microscale chambers separated from media chambers *via* an array of micropillars [24, 25]. These models have been utilized to study tumor-vasculature interactions [26], trafficking of nanoparticles across the blood-brain-barrier [27], and cell migration for prediction of survival and recurrence of GBM in patients [28]. Nevertheless, they fail to simulate pathophysiological features of tumors like compact cell-cell adhesion and diffusion gradient that affect the physiology of the cancerous cells [29].

The recent discovery of synaptic input of neurons on glioma cells [30], via a communication channel connecting neurons to glioma cells that affect glioma cell progression, [31] opened a novel clinical concept on how neuron-brain tumor synaptic interaction hampers tumor treatment [32, 33]. This finding emphasizes the necessity of direct interaction between neurons and glioblastoma cells for recapitulating glioblastoma microenvironment. However, none of the previous microfluidic models have applied neurons to study GBM invasion and resistance.

Herein, we present a hybrid chemoresistant GBM model on-a-chip (HGoC) by hybridizing spheroid and microfluidic models to take advantage of both models and overcome the drawbacks of each. To this end, we produced tumor spheroids from TMZ-resistant glioma cells and co-cultured them with human-based neurons. We carried out proteomics analysis to compare TMZ-resistant and non-resistant glioma cells, cultured under 2D or 3D conditions, with human primary and recurrent glioblastoma samples. Our results revealed that resistant cells cultured in the form of spheroids (3D R) have the highest similarity with human recurrent GBM. Using lipidomics assay, we confirmed that our TMZ-resistant engineered tissues recapitulate the molecular signature of human recurrent glioblastoma. We further characterized the model using secretome assays that emphasized the significant effects of direct interaction with healthy surrounding neurons on the invasiveness and migration plasticity of resistant cells. Overall, our multi-omics approach proved the capability of HGoC in simulating the pathophysiology of human recurrent glioblastoma, verifying that HGoC can be used to investigate the mechanism of chemoresistance and to seek new therapeutic approaches.

## Results

### Hybrid TMZ-resistance Glioblastoma-on-a-chip (HGoC) model

This study used a multi-compartment microfluidic chip to model the GBM microenvironment. The chip comprised a middle chamber where the tumor spheroids and neurons reside and two side channels for delivering nutrients (Fig. 1a). A composite hydrogel (CH) containing matrigel and alginate was used to mimic the extracellular matrix (ECM) of the brain. Matrigel is a natural murine basement membrane matrix that meticulously replicates the ECM components of the brain [34–36]. Alginate is a natural polysaccharide that mirrors ECM structure and can be tuned to adjust the stiffness of the matrix [37, 38]. We selected a composition of Matrigel and alginate based on an RNA-seq study reported the best to recapitulate the human brain ECM [39]. Bright-field and confocal images of HGoC showed that HGoC enabled the formation of direct interaction between glioma cells and surrounding neurons and invasion of R cells to the surrounding neurons, respectively (Fig. 1f, g).

**Fig 1.**
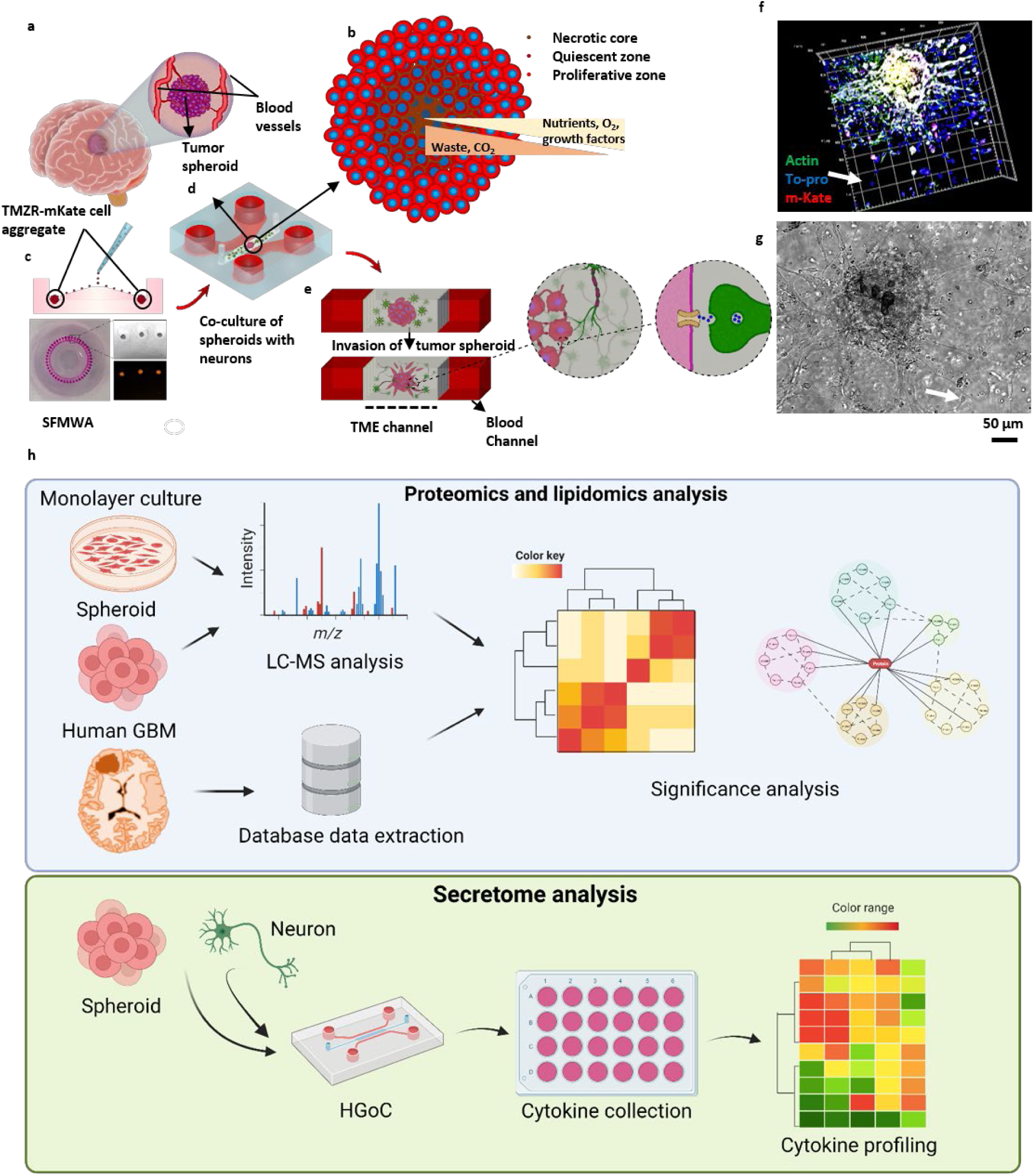
Hybrid microphysiological model of chemoresistant glioblastoma-on-a-chip (HGoC). a) A schematic of a brain tumour and surrounding capillaries. b) A schematic of tumor spheroid, showing the decreasing gradient for nutrient toward the core of spheroid and increasing gradient of waste out of the spheroid. c) Formation of chemoresistant spheroids on SFMWA. d) A schematic of HGoC designed to mimic brain tumor niche in tumour microenvironment channel and blood channels. e) Invasion of tumor spheroids to surrounding tissues. Magnified picture shows the direct interaction of GBM cancerous cells with neuron and their potential synaptic interaction. f) Bright filed and CIF imaging of tumor spheroid invaded to surrounding neurons in TEM channel, blue: nucleus (neurons), green: actin filament, red: mKate protein, white: merged (mKate cells). h) workflow of the study. The top panel is showing the steps on recapitulating resistant human glioblastoma and characterization of resistant spheroids. The bottom panel is showing the steps on recapitulating tumor microenvironment and <u>characterizing</u> the model. R: resistant, NR: non-resistant, MWA: Microwell arrays, TME: Tumor microenvironment, CIF: Confocal immunofluorescent, SFMWA: self-filling microwell arrays

To model recurrent GBM, we first established a TMZ-resistant glioma cell line using a pulsed-selection strategy [40]. This method led to the production of TMZ-resistant U251 cells with a half-maximal effective concentration (EC_50_) of 674 µM, which was six times higher than the non-resistant cells (111 µM) (Supplementary Fig. 1a, b). To investigate the chemosensitivity of the established cell lines in 3D, non-resistant and TMZ-resistant tumor spheroids (200 µm in diameter) were formed using our previously developed self-filling microwell array (SFMWA) (Supplementary Fig. 1c)[41]. TMZ-resistant spheroids (3D R) displayed significantly lower sensitivity to TMZ (5000 µM) as compared with non-resistant spheroids (3D NR) (Supplementary Fig. 1d). Workflow showing the characterized the TMZ-resistant tumor spheroid using proteomic and lipidomic analysis and then the tumor microenvironment by secretome assay is displayed in Fig. 1h.

### HGoC show distinguished metabolic signature of the chemoresitant GB

To assess the diffusion of nutrients/metabolites, diffusion of FITC-dextran 20kDa was semi-quantified across the width of the TME channel in HGoC, as the cytokines and trophic factors in cerebral spinal fluid have a molecular weight between 11-38kDa [42]. TME channel of HGoC is diffusible for 20kDa chemicals as fast as 12.5 µm/min (Fig. 2a, b). CH shows a biocompatibility by more than 70% viable U251-mKate cells and 50% viable neurons after five days in HGoC (Fig. 2c, d). Confocal immunofluorescence (CIF) imaging of neuronal markers, Map-2 and Tuj1 [8], confirms the successful differentiation of NPC cells toward neurons (Fig. 2e).

**Figure 2.**
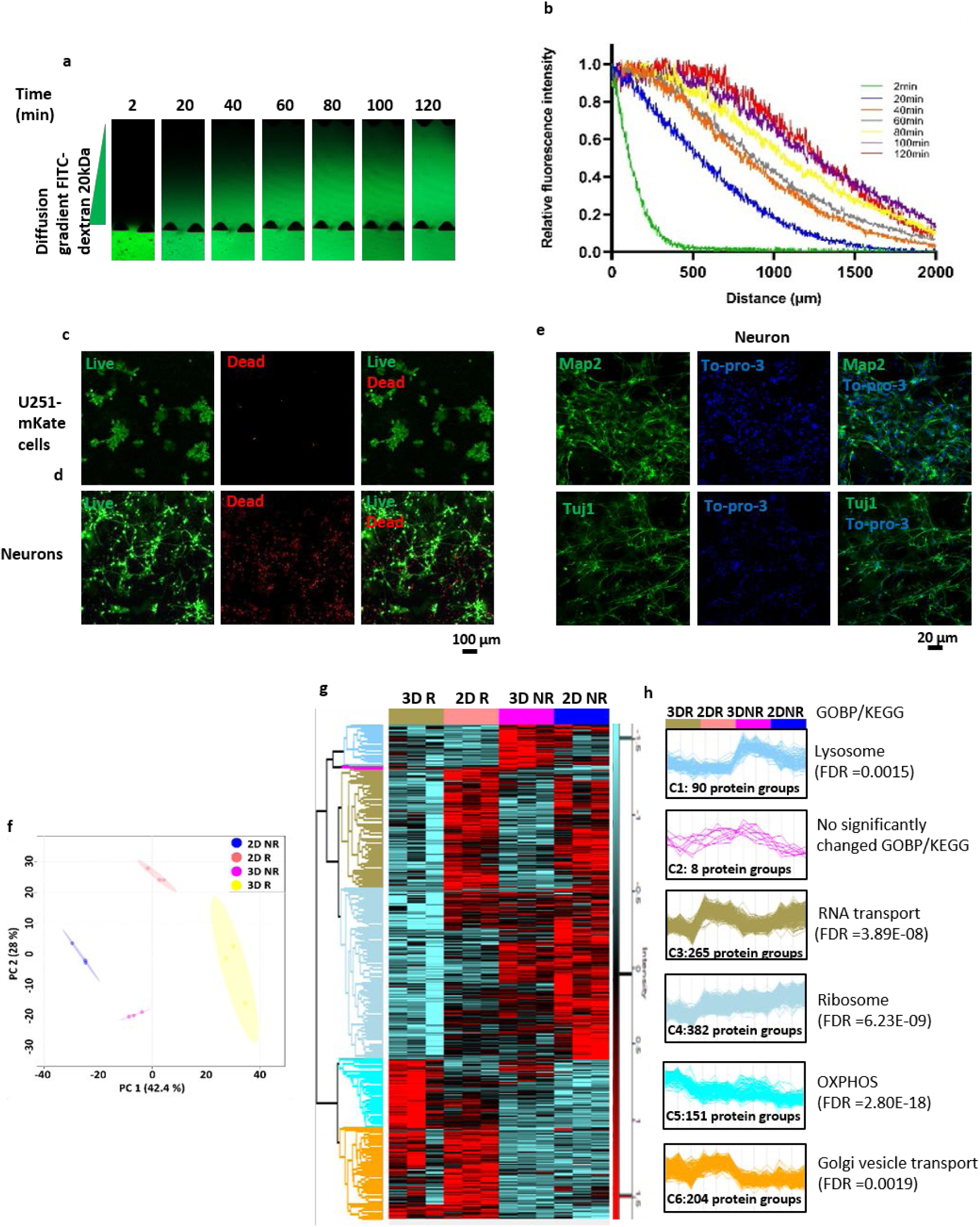
Global characterization of HGoC. a) Diffusion of FITC-dextran 20kDa from blood channel into the TME channel through the composite hydrogel. b) Quantitative analysis of fluorescent intensity of diffused FITC-dextran 20kDa. c) Live/dead assay of U251-mKate cells after 4 days encapsulation in CH and injection inside the TME channels. Scale bar=100 μm. d) Live/dead assay of neuron after 5 days encapsulation in CH and injection inside the TME channels. f) Principal component analysis demonstrates a clear distinction between the proteomes of 2D, 3D, R and NR samples. e) CIF imaging of Map2 and Tuj1, markers for neurons, and To-pro-3, marker of nucleus. Scale bar=50 μm. g) Hierarchical clustering of normalized protein concentrations. Each raw represent a distinct protein and each column represent a sample. 2D R: resistant glioblastoma cells cultured in 2D, 2D NR: non-resistant glioblastoma cells cultured in 2D, 3D NR: non-resistant glioblastoma cells cultured in spheroid form, 3D R: resistant glioblastoma cells cultured in spheroid form (one-way ANOVA: Benjamini–Hochberg FDR = 0.05. h) Protein expression profiles for each cluster and the most enriched KEGG/GOBP per cluster is shown on the right side of each profile. N=3 biological independent experiment in (a-e & g). CIF: Confocal immunofleurescent, FITC: Fluorescein Isothiocyanate, TME: Tumor microenvironment.

Principal component analysis was performed on all quantified proteins for an insightful overview of the biological bases of acquired resistance and the effect of culturing in 3D (Fig. 2f). Conspicuously, acquired resistance differentiates 2D R from 2D NR with higher expression of variables on both PC 1 and PC 2. However, culturing in 3D provided a unique feature, separating 3D R from 2D R with higher expression of variables on PC 1 but lower expression of variables on PC 2. Although 3D R and 3D NR indicate similarities on PC 2, a clear distinction on PC 1 was observed. To ensure our proteome data’s reproducibility, we estimated the correlation between every two samples by the Row-wise method and depicted the correlation using the Scatterplot Matrix. We found a considerable direct correlation between all of the two samples (0.85 ≥ R ≤ 1.0; Supplementary Fig. 2).

Hierarchical clustering found 1100 differentially regulated proteins in response to acquiring resistance and culturing in 3D and determined the direction of regulation of proteins (Fig. 2g). Enrichment analysis identified six expression clusters and showed that acquiring resistance is associated with decreased expression of lysosomal and ribosomal proteins. In contrast, it is associated with increased expression of genes involved in oxidative phosphorylation (OXPHOS) and vesicular transport from and to Golgi (Fig. 2h, C1,4-6, Supplementary Table 1.). In addition, it reveals that culturing under 3D conditions is associated with lower expression of proteins involved in RNA transport (Fig. 2h, C3, Supplementary Table 1.).

### Proteome signature of acquired resistance: mesenchymal stemness, mitochondrial respiration, vesicular transport, and mitotic quiescent

To gain insights into the molecular bases of the acquired resistance, we performed a volcano plot analysis to compare resistant and non-resistant cells in 2D and 3D cultures (Fig. 3a & b, Supplementary Table. 2). Most of the proteins that are upregulated more than two folds in resistant cells and spheroids compared with the non-resistant ones are known to be associated with decreased programmed cell death (BST2 [43], FTL and FTH1 [44], ASAH1 [45]), the induction of mesenchymal stemness (ALDH2 [46], SERPINH1 [47]), or arrest of the cell cycle (3IFITM [48], ISG15 [49, 50], p53 [51]). On the other hand, among the proteins that are downregulated more than twice, glial fibrillary acidic protein (GFAP) is the marker of glial cell differentiation [52], another finding that suggests the stemness character of resistant cells. Interestingly, the expression of GFAP has decreased eleven times in 3D R compared to 3D NR. However in 2D condition this decrease was three times, suggesting that the 3D condition enhances the stemness of cells [53]. Cathepsin L1 (CTSL), a lysosomal cysteine peptidase, is decreased nine folds in 3D R compared to 3D NR, while this decrease is less than two times in 2D condition (Fig 2h). This visualize further lower expression of lysosmal proteins in 2D conditions compared with 3D suggested by the enriched KEGG pathway in cluster 1 (Fig. 2h, C1).

**Fig 3.**
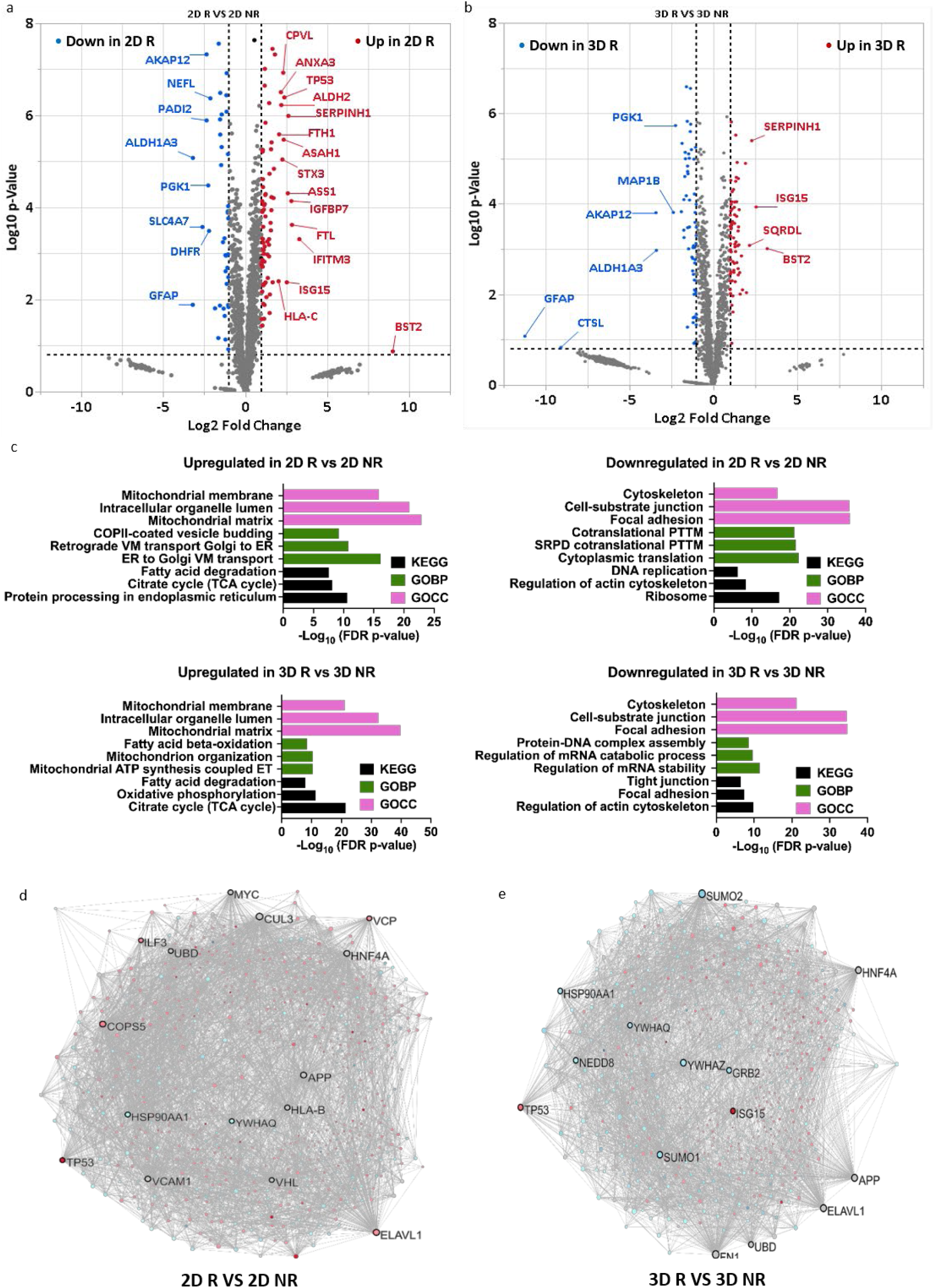
Molecular signature of acquired resistance. a) Volcano scatter plot of non-resistant vs resistant glioblastoma cells cultured in 2D (2D NR vs 2D R). The red or blue dots indicate proteins significantly (q < 0.05) upregulated (log_2_ fold change ≥ 0.5) or downregulated (log_2_ fold change ≤ −0.5) respectively. Proteins with more than 2 fold up or downregulation are annotated. b) Volcano scatter plot of non-resistant vs resistant glioblastoma cells cultured in 3D (3D NR VS 3D R). The red or blue dots indicate proteins significantly (q < 0.05) upregulated (log_2_ fold change ≥ 0.5) or downregulated (log_2_ fold change ≤ −0.5) respectively. Proteins with more than 2 fold up or downregulation are labeled. c) The three most significant GO terms and KEGG pathways that up/down regulated in each group emerged from the enrichment analysis of the genes identified by student t-test. d) Protein-protein interaction networks were constructed using Network Analyst. Nodes up-regulated in 2D R are shown in red and nodes down-regulated in 2D R are shown in blue and first order interactions are shown in grey. e) Protein-protein interaction networks were constructed using Network Analyst. Nodes up-regulated in 3D R are shown in red and nodes down-regulated in 3D R are shown in blue and first order interactions are shown in grey. ATP: Adenosine triphosphate. ER: endoplasmic reticulum. ET: electron transport. PTTM: protein targeting to membrane. SRPD: signal-recognition particle (SRP)-dependent. TCA: tricarboxylic acid. VM: vesicular mediated.

To define enriched gene ontology terms and functional pathways, enrichment analysis was executed on up- or downregulated proteins using the Enrichr database (Fig 3c) [54–56]. The citric acid (TCA) cycle and fatty acid degradation are the most enriched KEGG pathways in both conditions. In line with upregulated pathways, mitochondrial structures are the most enriched cellular components. Enrichment in both KEGG pathways, fatty acid degradation and citrate cycle, and cellular components, mitochondrial matrix and membrane in 3D condition are twice significant than in 2D condition that emphasize the occurred changes. Regarding GOBP, in 2D conditions, anterograde and retrograde vesicular mediated transport between Golgi and endoplasmic reticulum (ER) were the most enriched upregulated biological processes. In contrast, in 3D conditions, ATP synthesis *via* the electron transport chain and fatty acid beta-oxidation in mitochondria were the most enriched upregulated biological processes.

Enriched analysis of downregulated gene ontology terms (Fig 3c) shows that regulation of actin cytoskeleton is among the three top most enriched downregulated KEGG pathways, and synchronously cytoskeleton is the most enriched downregulated cellular component in both 2D and 3D conditions. Focal adhesion and tight junctions are the other most enriched downregulated pathways in 3D conditions, while DNA replication and ribosomes are the most enriched downregulated pathways in 2D conditions. Other enriched downregulated cellular components show decreased mechanical links between intracellular actin and ECM through focal adhesions and cell-substrate junctions in 2D and 3D conditions. The most enriched downregulated biological processes in 2D and 3D conditions show decreased protein translation.

To understand data at the system level, protein-protein interactions (PPI) were constructed to create 2D R-specific (Fig. 3d) and 3D R-specific PPI networks (Fig. 3e). The top upregulated Reactome pathway associated with top nodes is TCA Cycle and Respiratory Electron Transport in both 2D R (FDR=1.00E-08) and 3D R (FDR=4.34E-30) (Supplementary Table 3). The top downregulated Reactome pathway associated with top nodes displays decreased translation and apoptosis in 2D R (FDR=1.00E-06) and 3D R (FDR=4.47E-06), respectively (Supplementary Table 3). Top upregulated GOBP associated with top nodes reveals enhanced vesicular transport between Golgi-ER and cell respiration in 2D R (FDR=5.18E-14) and 3D R (FDR=6.28E-27), respectively (Supplementary Table 3). Top downregulated GOBP associated with top nodes displays decreased translation and mitotic cell cycle in 2D R (FDR=5.17E-11) and 3D R (FDR=1.29E-08), respectively (Supplementary Table 3).

Overall, acquiring resistance was associated with increased expression of genes involved in DNA repair and cell survival and upregulation of vesicular transport, TCA cycle, and OXPHOS, a metabolic state essential to sustain mesenchymal stemness character [57]. In addition, resistant cells showed a decreased expression of genes involved in cell differentiation and downregulation of cell-cell and cell-substrate junctions and cytoskeleton.

### TMZ-resistance was associated with epithelial-to-mesenchymal transition (EMT), mitotic quiescence and OXPHOS metabolism

The EMT, a transition from a differentiated to an undifferentiated state, is frequently reported in cancer resistance, progression, and metastasis [58–61]. To analyze EMT in glioma cells, the morphology of resistant and non-resistant cells was inspected using bright-field microscopy. Interestingly, resistance cells displayed significant morphological changes featured by elongated cells (Fig. 4a) with a high amount of cytoplasmic vesicles Supplementary Fig 3d) —both characteristics of mesenchymal phenotype [61]. Immunoblotting further confirmed our observations by upregulation of mesenchymal markers, N-cadherin and Vimentin, in resistant cells (Fig. 4b).

**Fig 4.**
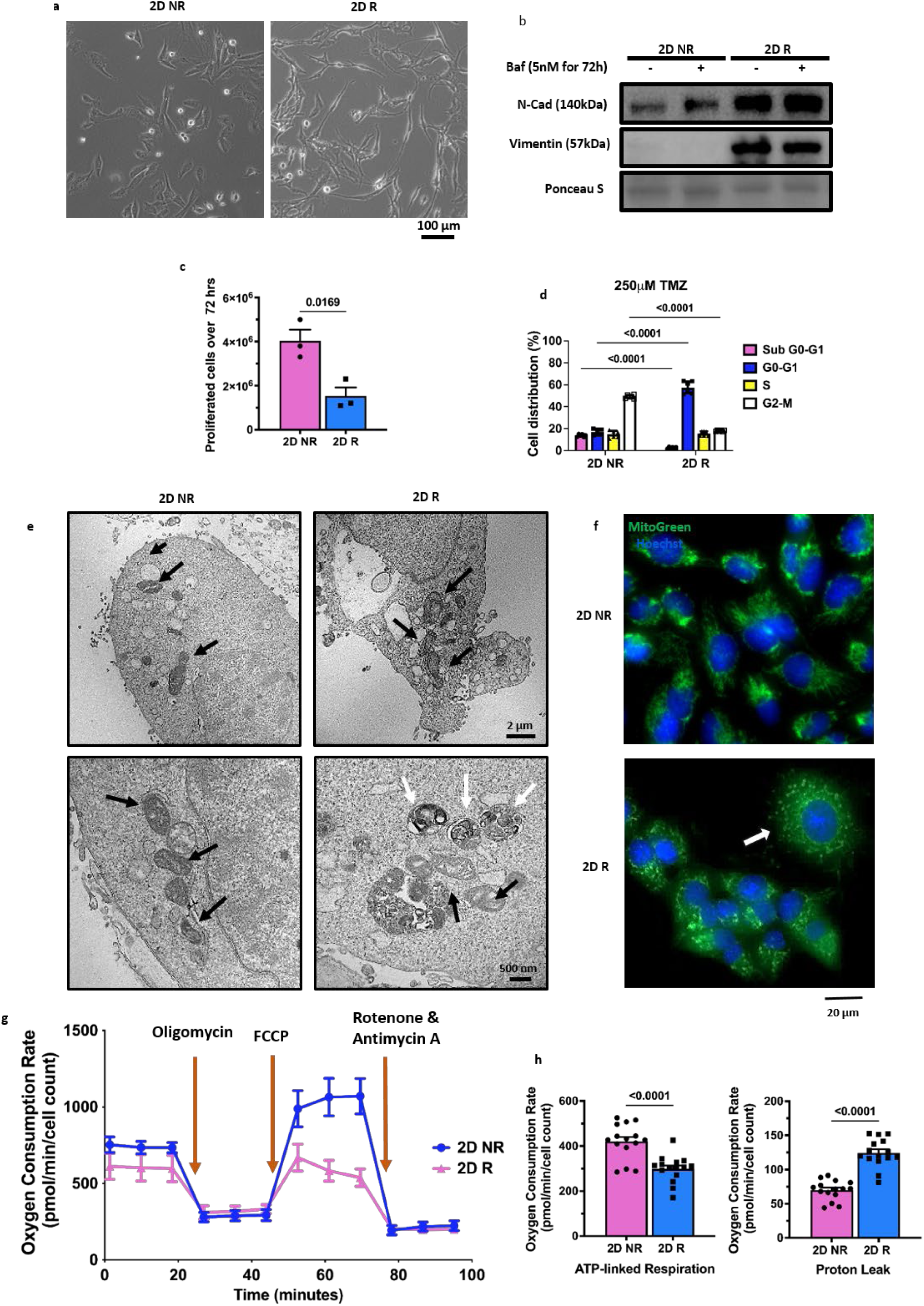
Stem-like signature of acquired resistance. a) Bright field imaging comparing 2D R with 2D NR. b) Western blotting of N-cadherin and Vimentin compared between 2D NR and 2D R without treatment and after treatment with Bafilomycin a1 5nM for 72hrs. c) Proliferation assay comparing the number of the cells 72hrs after seeding 500 cell/ml. d) Flow cytometer results of cell cycle assay comparing 2D R with 2D NR after 48hrs treatment with 250µM TMZ. e) TEM images compared mitochondrial structure between 2D NR with 2D R. Black arrows show the mitochondria and white arrows show the mitophagosomes. f) Live cell IF imaging of mitochondria using mitoviewGreen, arrow shows mitochondria. g) Oxygen consumption rates (OCR) were analyzed using a Seahorse XFe24 Analyzer and normalized to cell count. h) Mean ATP production, and proton leak normalized to cell count were calculated. Scale bar=20 um. TEM: The transmission electron microscope. N=3 and 8 biological independent experiment in (a-f) and (g-h) respectively.

Autophagy has been introduced as a mechanism of chemoresitance in cancers [62, 63] including glioblastoma [14, 64]. Regarding the pivotal role of autophagy in the regulation of EMT [65, 66], we also studied the effect of autophagy flux inhibitor on the expression of mesenchymal markers that showed treatment with Bafilomycin A1 (Baf) 5 nM for 72 hrs did not affect the expression of these markers in non-resistant and resistant cells (Fig. 4b).

To study the mitotic quiescence, another feature of stem cells [6], we analyzed the proliferation rate and cell cycle state of resistance with non-resistant cells. The number of cells after 72 hrs decreased significantly in resistant cells, and the proliferation rate to less than half of non-resistant cells (Fig. 4c). Cell cycle analysis by flow cytometry revealed that the population of mitotic cells decreased more than two folds in resistant cells, while the population of cells arrested in G1 increased three folds, compared to non-resistant cells. Our results also showed a drastic drop in the population of apoptotic cells (sub G0-G1) (Fig. 4d).

Switch of metabolism in cancer stem cells has been shown as a mechanism of adaptation [58, 67]. To confirm the proteomics results showing increased mitochondrial oxidation of TCA and fatty acids, we performed transmission electron microscopy (TEM) imaging that demonstrates significant structural changes in mitochondria associated with resistance, including abnormal cristae and increased mitophagy (Fig. 4e). Live cell immunofluorescence imaging of mitochondria using MitoView green, which relied on mitochondrial mass, presents elongated mitochondria phenotype in non-resistant cells. In contrast, we observed mitochondrial fragmentation and clustering in resistant cells, suggesting mitochondrial fission and mitophagy [68], consistent with the TEM results on mitophagy (Fig. 4f). Mitophagy is critical in acquiring stemness and metabolic switch to OXPHOS [68]. Measurement of mitochondrial respiration indicated lower basal mitochondrial respiration and ATP-linked respiration in resistant cells compared to the non-resistant cells due to the lower proliferation rate and metabolic activity in resistant cells (Fig. 4g & h). However, resistance was associated with higher proton leak, a mechanism that regulates the mitochondrial ATP production (Fig. 4g & h).

Altogether, these data show that acquiring resistance is associated with EMT in 2D and MAT in 3D. In addition, acquiring resistance resulted in mitotic quiescent and a switch of metabolism toward the TCA cycle and OXPHOS.

### TMZ-resistance was associated with inhibition of autophagy flux

We performed TEM imaging to validate the upregulated vesicular trafficking suggested by proteomics results (Fig 3c). TEM results showed an increased vesicular structure associated with acquiring resistance and a higher number of double-membrane vesicles that are the autophagosome criteria (Fig. 5a & b). Live cell IF imaging of LC3, the marker of autophagosome formation, showed an increase in the number of autophagosome vesicles in 2D R compared with the 2D NR (Fig. 5c). To determine whether this finding was due to upregulation of the autophagy pathway or inhibition of autophagy flux, we performed western blotting of LC3 and P62, the marker of autophagosome degradation (Fig. 5d). Our results showed that inhibiting autophagy flux using Baf, which suppresses lysosomal acidification, caused no changes in the level of autophagy markers (Fig. 5e). However, live cell imaging with Lysotracker green showed no changes in acidic compartments between resistant and non-resistant cells (Fig. 5f). These findings suggest that impairment in autophagosome maturation to autolysosome is the mechanism of autophagy flux inhibition, which is in line with our findings from hierarchical clustering that showed decreased lysosomal protein expression in resistant cells (Fig. 2h). Overall, we speculate that TMZ resistance was associated with increased vesicular trafficking, which is partly related to the inhibition of autophagy flux.

**Fig 5.**
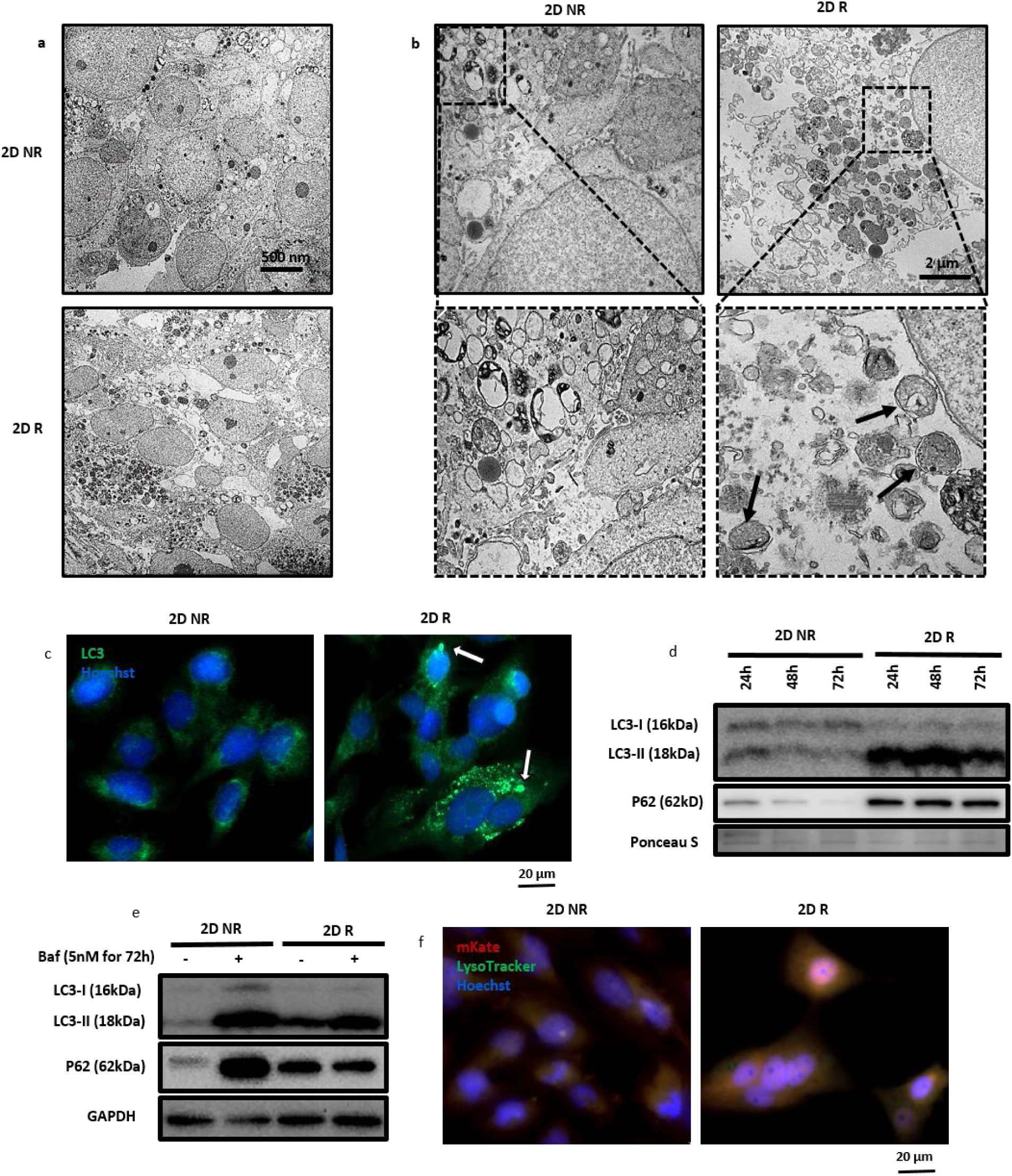
Enhanced vesicular transport in acquired resistance. a & b) TEM images compared cellular and intracellular structure between 2D NR with 2D R, arrows show the autophagosomes. c) Live cell IF imaging of LC3, white arrows show LC3 punctate, representative of autophagosome formation. d) Western blotting of LC3 and P62 compared between 2D NR and 2D R. Ponceau S staining confirm homogeneous protein loading and transfer. e) Western blotting of LC3 and P62 compared between 2D NR and 2D R with and without 72hrs treatment with Baf 5nM. GAPDH confirm homogeneous protein loading and transfer, and to normalize the results. f) Live cell staining with LysoTracker green show no changes in cell acidic compartments between resistant and non-resistant groups. N=3 biological independent experiment in (a-f).

### Cells grown in 3D exhibited increased mitochondrial oxidation, high cell viability, decreased attachment properties and mesenchymal to amoeboid transition (MAT)

To elucidate the effect of 3D culturing on the phenotype of resistance cells, we conducted a volcano plot analysis, comparing proteomics data for 2D with 3D and highlighted only the proteins with the degree of regulation more/less than two folds. The results show that the expression of proteins such as GFAP, Cathepsin L (CTSL),, and Calcyphosin (CAPS) that were downregulated while acquiring the resistance, were upregulated in non-resistant cells cultured in 3D (Supplementary Fig. 3a & 3b, Supplementary Table 2). Enrichment analysis on up or downregulated proteins using the Enrichr database reveals that 3D condition resulted in enhanced mitochondrial oxidation primarily due to an increased TCA cycle and OXPHOS in resistant cells. In contrast, in non-resistant cells, fatty acid oxidation (FAO) was the most enriched KEGG pathway (Fig 6a). As expected from enriched KEGG pathways, the mitochondrial matrix was the most enriched cellular component in cells grown in 3D as compared to those grown in 2D. Fatty acid beta-oxidation is the most enriched biological process in 3D R and 3D NR. In contrast, aerobic electron transport and epithelial cell differentiation are exclusively enriched GOBP in 3D R and 3D NR, respectively.

**Fig 6.**
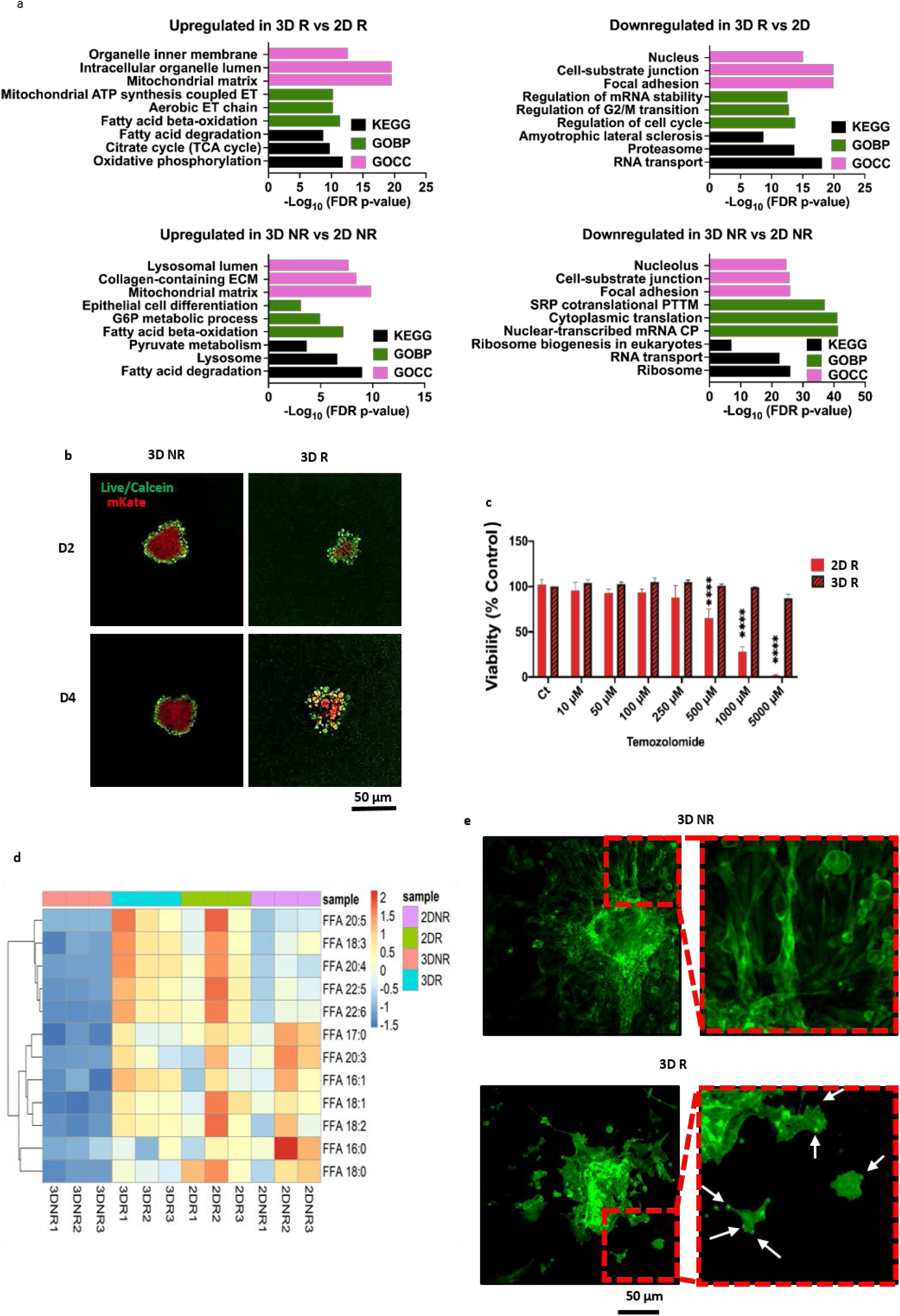
Molecular and cellular signature of culturing under 3D structure. a) Top three most significant GO terms and KEGG pathways that up/down regulated in each group emerged from the enrichment analysis of the genes identified by student t-test. b) Confocal imaging to compare viability of tumorospheres expressing red fluorescent protein mKate that stained with calcein (stains live cells in green) in day 2 and day 4 after spheroid formation. c) Viability assay comparing viability of resistant cell cultured in 2D with 3D using presto blue assay. d) Hierarchical analysis of free fatty acids (FFA). e) Confocal imaging of actin filaments, visualized with phaloidin staining, compared the invasion pattern of 3D R and 3D NR. d) Proliferation assay comparing cell number of resistant and non-resistant cells 72 hrs after seeding of the 500k cells. N=3 biological independent experiment in (a-e). The values are mean ± SEM of n =15 biologically independent samples examined over 3 independent experiments in (c). ATP: Adenosine triphosphate. CP: catabolic process. ECM: extra cellular matrix. ET: electron transport. PTTM: protein targeting to membrane. SRP: signal-recognition particle. TCA: tricarboxylic acid.

Enriched analysis of downregulated gene ontology terms showed that RNA transport and cell-substrate junctions were commonly enriched downregulated KEGG pathways and cellular components in 3D R and 3D NR, respectively (Fig 6a). The most enriched downregulated biological process in 3D R was related to cell mitosis, while in 3D NR, it was related to cytoplasmic translation. Accordingly, the nucleus and nucleolus are among the top three enriched cellular components in 3D R and 3D NR, respectively. Uniquely, amyotrophic lateral sclerosis, a pathway that resulted in neuronal cell death, was amongst the top three enriched downregulated KEGG pathways in 3D R, implying higher viability of resistant cells under 3D conditions. However, in 3D NR, ribosome and ribosomal biogenesis were among the top three enriched downregulated KEGG pathways that show differential expression of proteins dependent on the cell type.

To confirm proteomics results on the effect of 3D culturing, we performed confocal imaging of spheroids stained with Calcein, a dye that specifically penetrates live cells (Fig 6b, individual channels are shown in Supplementary Fig 4). The results disclose that 3D R had higher viability and lower attachment property compared with 3D NR. To further investigate the effect of 3D culturing on drug sensitivity we compared the viability of resistant cells with resistant spheroids (Fig 6c). Interestingly, the result reveal that even 5000 µM of TMZ had slight effect on the viability of resistant cells when they were cultured under 3D condition, while this dose killed almost all of them when they were cultured under 2D condition. We ran lipidomics analysis and compared the levels of different species of free fatty acids (FFA) between our groups that confirm the proteomics results and shows that level of FFA were drastically dropped in 3D NR (Fig 6d), although this decrease was significant only on FFA 20:4, 22:5 and 22:6 (FDR≤0.05).

To further study invasion patterns of resistant cells, we performed confocal imaging of actin filament and compared invasion patterns of resistant and non-resistant spheroids encapsulated in hydrogel (3D R and 3D NR) (Fig. 5e). Interestingly, spheroids made of resistant cells exhibited an utterly different invasion pattern than their non-resistant counterparts. The non-resistant cells developed a star-shaped invasion pattern with extended protrusions into the healthy areas of the matrix, resembling diffuse gliomas. On the other hand, TMZ-resistant cells exhibited a scattered tumor growth with dispersed protrusions into the surrounding matrix. A closer look at the invading cells discloses that the non-resistant cells were elongated with clustered actin, finger-like projections into the surrounding TME, and filopodia protrusions [69], seen in mesenchymal cells [70]. Conversely, resistant cells were rounded with bleb-like projections that are shallow and actin-rich, podosomes protrusions [69], seen in amoeboid cells [70] (Fig. 5e). This migration plasticity confirms decreased adhesion properties in resistant cells related to mesenchymal to amoeboid transition (MAT), one of the stem cell characteristics [71–73].

In conclusion, culturing under 3D conditions enhances oxidative metabolism primarily due to higher TCA cycle and OXPHOS metabolism in resistant cells and FAO in non-resistant cells. Additionally, the 3D condition causes lower replication and programmed cell death in resistant cells. Further, it results in decreased attachment properties in resistant and non-resistant cells, although this decrease was more prominent in resistant cells visualized by podosome protrusions and MAT.

### 3D structures make resistant cells resemble human glioblastoma

To validate the extent to which our model represents human glioblastoma, we compared the proteomic phenotype of our samples with paired human primary (HP) and human recurrent (HR) glioblastoma patient tissues deposited by Dekker and colleagues to the ProteomeXchange Consortium via the PRIDE partner repository [74]. We identified 1007 differentially regulated proteins and determined the direction of regulation of proteins by hierarchical clustering. Notably, we classified thirteen expression clusters from which clusters 2, 7, 8, 10, and 12 (highlighted in red) showed the highest protein expression pattern similarity between 3D R samples and human glioblastoma samples (Fig. 7a).

**Fig 7.**
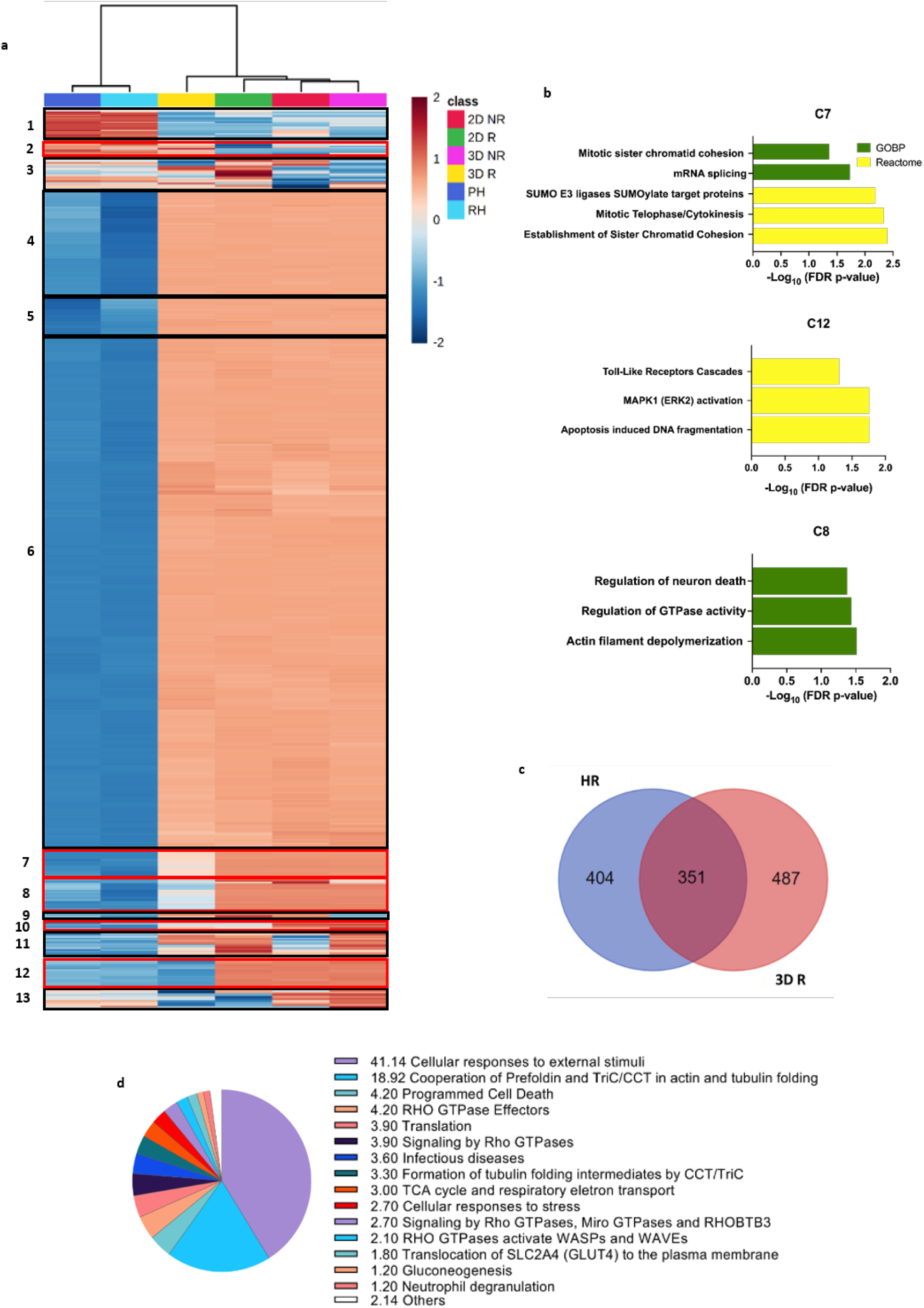
Comparing GBM engineered tissues with paired primary and recurrent glioblastoma patient tissues. a) Hierarchical clustering of normalized protein concentrations. Each raw represent a distinct protein and each column represent a sample. PH: human primary glioblastoma patients, HR: human recurrent glioblastoma patients, 2D R: resistant glioblastoma cells cultured in 2D, 2D NR: non-resistant glioblastoma cells cultured in 2D, 3D NR: non-resistant glioblastoma cells cultured in spheroid form, 3D R: resistant glioblastoma cells cultured in spheroid form (one-way ANOVA: Benjamini–Hochberg FDR = 0.05; n =3 biologically independent samples for 2D and 3D samples, n=8 for PH and RH). b) Top three enriched GOBP and Reactome pathways in clusters with similar protein expression pattern between human GBM tissues with 3DR. c) Venn diagram showing the number of differential and common proteome expression between HR and 3D R samples (adjusted *P* value is 0.05). d) Pie chart showing the functional classification of 351 proteins commonly expressed in both HR and 3D R.

Cluster 12, with the highest similarities between 3D R and human GBM patient tissues, demonstrates that culturing under three-dimensional conditions makes resistant cells resemble the proteome character of human GBM patients (Fig. 7a, C12). No gene ontology term was significantly enriched in this cluster. However, the most enriched Reactome pathways were MAPK1 activation and Toll-like receptors cascade involved in proliferation, differentiation, and adhesion (Fig. 7b). In addition, apoptosis-induced DNA fragmentation is another significantly enriched pathway down-regulated in both human GBM tissues and 3D R (Fig. 7b). Cluster 8 also represents a higher similarity in the proteomic character between 3D R and human glioblastoma than rest of the samples, especially with HR, which shows a lower expression of this cluster than HP (Fig. 7a, C8). No Reactome pathway is significantly enriched in this cluster. The most enriched gene ontology of the biological process is actin filament depolymerization, regulation of GTPase activity, and neuronal death, which are downregulated in both 3D R compared with our other tissues and HR compared with HP (Fig. 7a). Actin filament depolymerization is involved in stem cell differentiation [75, 76], and GTPase activity regulates the actin reorganization [77]. Thus, these data suggest dedifferentiation of our 3D R and HR toward stem cells. In cluster 7, the most enriched GOBP and Reactome pathways were related to translation and mitosis, both downregulated in 3D R and both human GBM tissues (Fig 7b). No gene ontology term or Reactome pathway were enriched in Cluster 2.

As expected, some clusters show differential expression of proteins between our samples and human tissues (Clusters 1, 4, 5, and 6), most likely due to the different nature of cell lines with primary cells [78]. Clusters 4, 5, and 6 display lower expression in human GBM tissues than in our engineered tissues. Nevertheless, they demonstrate similar expression patterns to our 3D R tissues. Proteins in cluster 4 with lower expression in HR than HP (Fig. 7a) are involved in replication, translation, and inflammatory responses, based on the most enriched GOBP and Reactome pathways (Supplementary Fig. 5). In addition, proteins in cluster 5 with higher expression in HR than HP are involved in OXPHOS metabolism and negative regulation of autophagosome assembly (Supplementary Fig. 5). These results reveal a similar pattern of protein translation between 3D R and HR on increased OXPHOS metabolism and decreased replication, translation, and inflammatory responses.

Interestingly, the most enriched GOBP and Reactome pathways in C1 with a higher expression in both HP and HR compared to all of our samples (Fig. 7a) are related to podosome assembly and folding of cytoskeleton proteins by chaperonin CCT/TRiC (Supplementary Fig. 5). These data suggest podosome protrusions of human GBM cells similar to our 3D R [69, 79]. Cluster 6 visualizes proteomes with differentially lower expression in both HP and HR compared to our samples while showing the same expression level in both HR and HP (Fig. 7a). The most enriched GOBP and Reactome pathways were mainly involved in the translational activity (Supplementary Fig. 5). The enrichment analysis of C13 with lower expression of proteins in HR and HP and our resistant tissues compared with our non-resistant tissues indicates decreased expression of proteins involved in carbohydrate metabolism and ECM organization (Supplementary Fig. 5).

A Venn diagram analysis showed that 351 proteins among 1242 proteins were commonly expressed in 3D R and HR (Fig. 7c). Functional classification of these commonly expressed proteins reveals that about 44% of proteins are involved in the cellular response to stimuli/stress, like a response to drugs. About 22% of proteins are involved in the actin-tubulin cell cytoskeleton, and 12% are involved in the Rho GTPase function that regulates actin cytoskeleton organization (Fig. 7d). The rest of the proteins are involved in the TCA cycle, respiratory electron transport, translation, and programmed cell death.

Overall, these results indicate that our engineered 3D R tissue simulated the proteomic characteristics of recurrent human glioblastomas like amoeboid phenotype, lower proliferation, translation, and apoptotic cell death (clusters 1, 12, and 7). They also revealed the upregulation of proteins involved in the TCA cycle and respiratory electron transport (cluster 5) and downregulation of proteins involved in replication and translation in HR compared to the HP (clusters 4 and 8). A similar pattern was observed in 3D R compared with our other groups.

### Engineered resistant glioblastoma model simulates the hallmark of human recurrent glioblastoma

Dekker and colleagues, in a multi-omics study comparing paired primary and recurrent glioblastoma patient tissues, reported the cholesterol pathway as one of the pathways mostly affected at different omics levels when comparing cells collected from recurrent and prima glioblastoma patients [74]. Interestingly, cluster 10 had shown a similar pattern of decreased expression in human glioblastoma and our resistance samples (Fig. 8a). Reactome analysis displayed the cholesterol biosynthesis pathway assignificantly affected, and the most enriched GOBP is the Isopentenyl Diphosphate biosynthetic process, a pivotal cholesterol pathway component (Fig. 8b).

**Fig 8.**
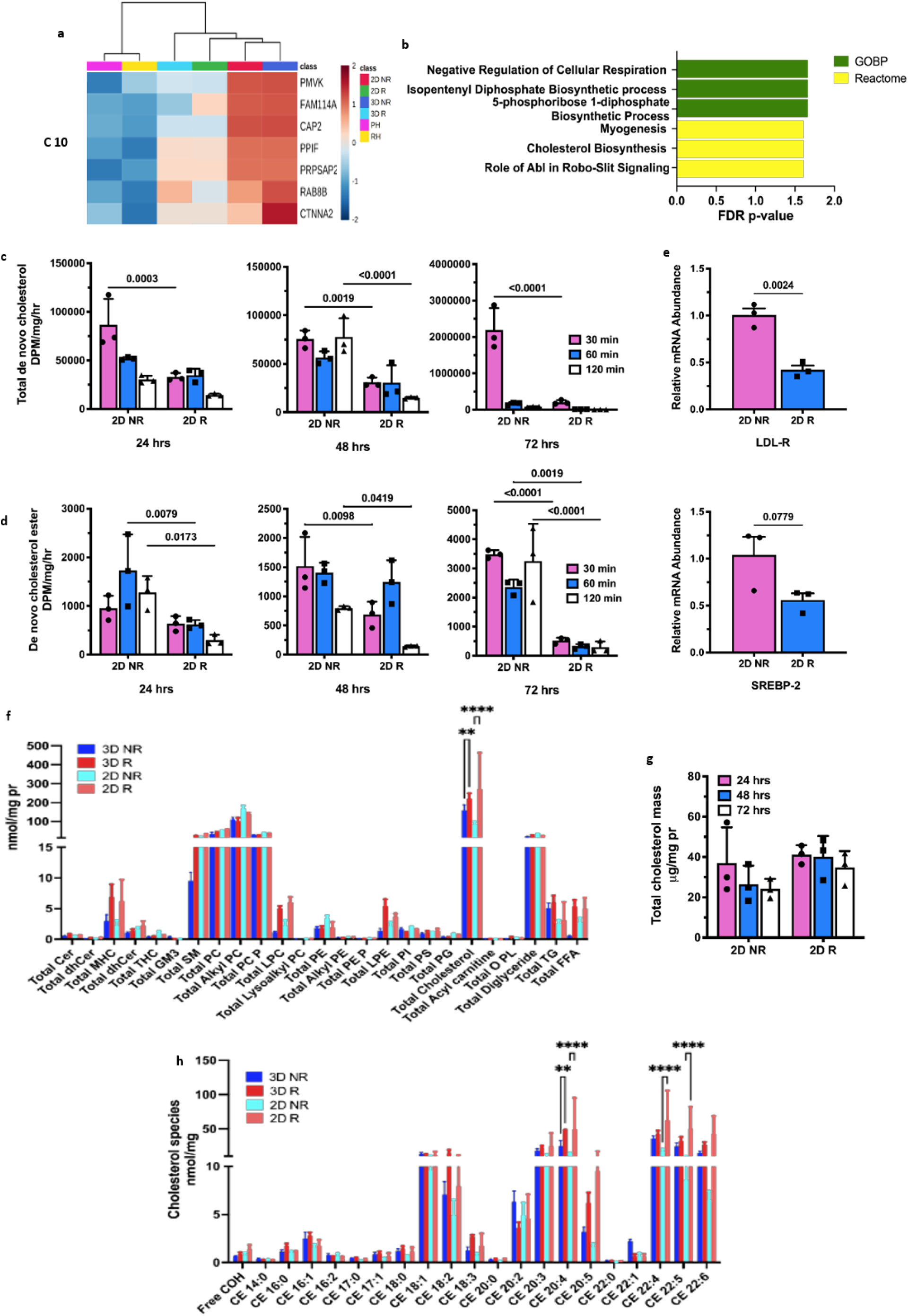
Acquiring resistance is associated with decreased cholesterol biosynthesis. a) Cluster 10 of heatmap. b) Top GOBP and Reactome pathways significantly affected in C10. c) Total de novo synthesized cholesterol after 24, 48 and 72 hrs culture. It revealed significant decrease in total de novo synthesized cholesterol in 2D R cells in all timepoints. d) De novo Cholesterol esterification after 24, 48 and 72 hrs culture. e) Real-time PCR measured the expression level of LDL-R and SREBP-2 genes. f) Lipidomics analysis of total lipid species. g) Total cholesterol mass measurement between resistant and non-resistant groups. h) Lipidomic analysis of cholesterol species. N=3 biological independent experiment in (c-h).

To confirm the proteomics results, we compared the total *de novo* synthesis and *de novo* esterification of cholesterol between R and NR samples (Fig. 8c & d). Both *de novo* synthesis and *de novo* esterification of cholesterol in all time points are decremented in 2D R compared with 2D NR. To elucidate if this is reflected at the transcription level, we performed real-time PCR. We compared the mRNA level of LDL-R (low-density lipoprotein receptor), a receptor for LDL that transports cholesterol into the cells, and SREBP2 (Sterol regulatory-element binding proteins), a transcription factor that regulates cholesterol homeostasis [80], between R and NR samples (Fig. 8e). A significant reduction in the mRNA level of LDL-R and SREBP2 in resistant cells compared with the non-resistant cells confirms the findings from proteomics and *de novo* cholesterol assays.

To further explore the role of lipid metabolism in glioblastoma resistance, we conducted a lipidomic analysis. The lipid profile of the total amount of different lipid species shows cholesterol as the only lipid species significantly affected between resistant and non-resistant groups (Fig. 8f). This finding contrasted with the results from *de novo* biosynthesis of cholesterol (Fig. 8c). Thus, we evaluated the total cholesterol mass that reveals higher total cholesterol mass in R cells than in NR cells, although this elevation is not significant (Fig. 8g). This finding suggests a decrease in the degradation of cholesterol molecules that resulted in their accumulation despite the decrease in cholesterol production. To further examine this hypothesis, we defined the cholesteryl ester profile, which demonstrated a higher amount of cholesteryl esters with a high degree of unsaturation, 20:4, 22:4, and 22:5, in the R groups compared to NR ones (Fig. 8h), cholesteryl esters that their degradations depend on lysosomal acid lipase [81].

Overall, the proteomic results show that in our 3D R tissue, cholesterol pathway was significantly affected, mimicking the hallmark of human recurrent glioblastoma. This finding was confirmed with lipidomic analysis and measurement of *de novo* cholesterol biosynthesis and esterification.

### HGoC recapitulates microphysiology and micropathophysiology of recurrent GBM tumors and its microenvironment

Considering the pivotal role of tumor microenviroment including ECM and cells in tumor physiology [18, 82], especially in GBM due to a synaptic integration between glioma cells and neurons [30, 31], we directly co-cultured our GBM tissues (3D R and 3D NR) with neurons and encapsulated them in a composite hydrogel comprised of alginate and Matrigel (HGoC and NHGoC). The invasive phenotype of HGoC with NHGoC were monitored for up to 4 days. Spheroid diameter and radius of invasion were quantified as the two parameters related to cell proliferation, invasion, and migration [83, 84] (Fig. 9b). We observed a star-shaped invasion in NHGoC and scattered invasion in HGoC (Fig. 9a, Fig. 4c). Further, spheroids in HGoC grew at a much slower rates when compared to spheroids in NHGoC (Fig. 9b).

**Fig 9.**
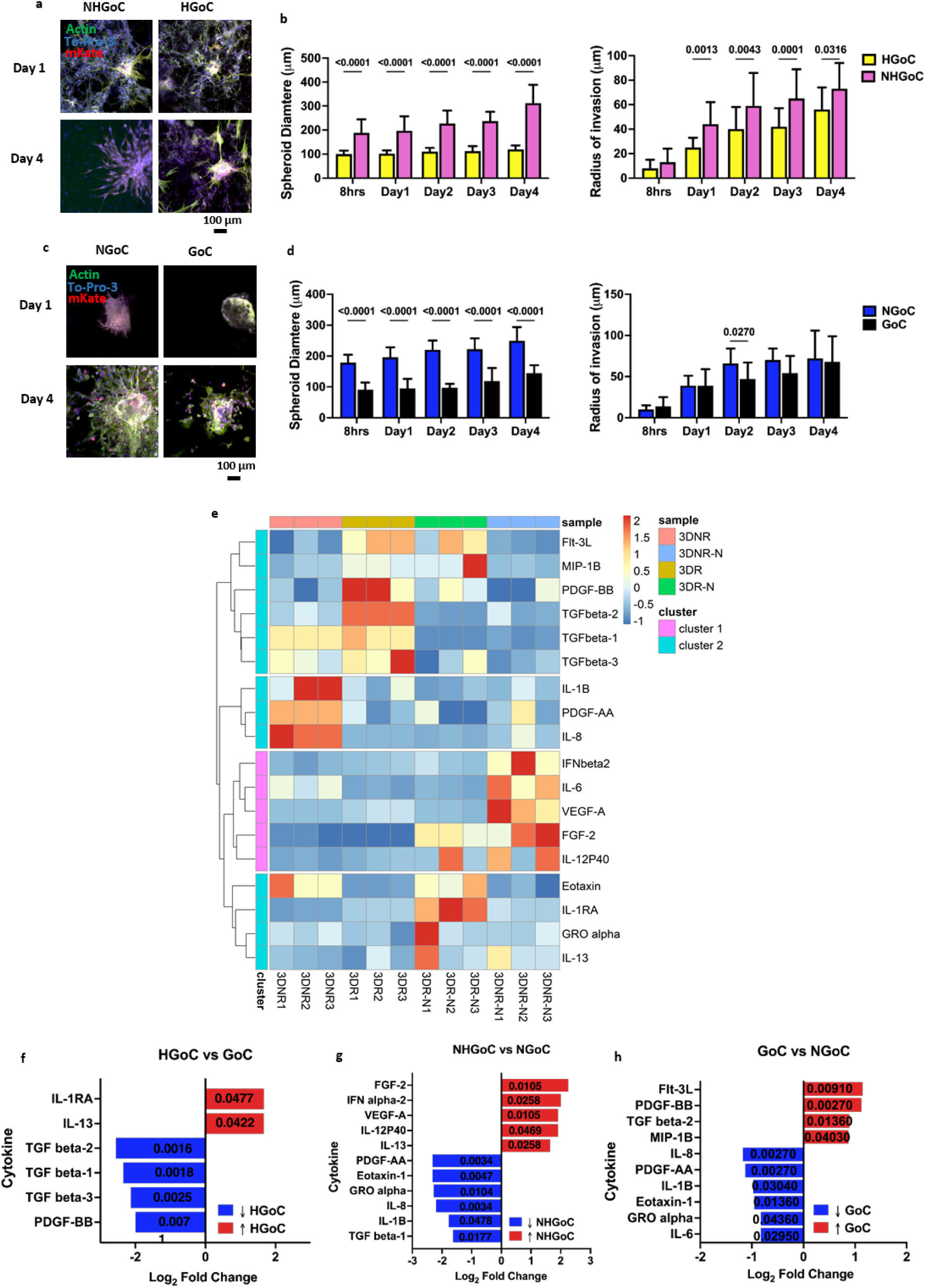
Direct interaction with neurons decrease invasiveness and immunomodulation property of resistance cells. a) Confocal imaging comparing day 1 and day 4 of NHGoC with HGoC, which were stained with phaloidin to visualize actin filament and to-pro3 to visualize nucleus. b) Quantitative analysis of diameter of tumorospheres and radius of invasion of NHGoC vs HGoC form 8hrs after injection inside the chip until day 4 of culture using imageJ software using imageJ software. c) Confocal imaging comparing day 1 and day 4 of NGoC with GoC, which were stained with phaloidin to visualize actin filament and to-pro3 to visualize nucleus. d) Quantitative analysis of diameter of tumor spheroids and radius of invasion of NGoC VS GoC form 8hrs after injection inside the chip until day 4 of culture using imageJ software. e) Hierarchical clustering of normalized cytokine concentrations. Each raw represent a distinct cytokine and each column represent a sample (significant cytokine was extracted based on the difference in their expression values with log2 fold change ≥ 2, FDR = 0.001). f) Cytokine profile comparing 3D R with 3D NR. g) Cytokine profile comparing NGoC with NHGoC. h) Cytokine profile comparing GoC with HGoC. NGoC: Non-resistant spheorids (3D NR)-without neurons. GoC: Resistant spheorids (3D R)-without neurons. NHGoC: Non-resistant spheorids (3D NR)-with neurons. HGoC: Resistant spheorids (3D)-with neurons. N=3 biological independent experiment in (a-h).

To study the impact of neuronal interaction on GBM spheroids’ phenotype and behavior, we compared the invasive profile of GoC (3D R encapsulated in composite hydrogel) with NGoC (3D NR encapsulated in composite hydrogel) (Fig. 9c). Although the increase in the spheroid diameter in NGoC was higher than in GoC, this increase was lower in the absence of neurons in non-resistant spheroids compared with resistant ones, 300 µm in NHGoC to 200 µm in NGoC on day 4. Yet, the spheroid diameter remains almost the same (≈100 µm) in both HGoC and GoC (Fig. 9 b & d). In addition, almost no significant changes are observed in the radius of invasion between GoC and NGoC (Fig. 9d). Radius of invasion decreased from ≈70 µm in GoC to < 60 µm in HGoC on day 4, remaining the same ≈ at 70 µm in both NHGoC and NGoC on day 4 (Fig. 9 b & d) suggests decreased invasiveness of resistant spheroids in the presence of neurons.

To gain an insight into the effect of neurons on the function of the engineered glioblastoma, we obtained the secretome profile of 68 cytokines from the inflammatory/invasion family in our engineered tissues, among which 18 cytokines were significantly altered (Fig. 9e). A separate cluster in the cytokine heatmap for NHGoC displayed that the presence of neurons led to the higher secretion of cytokines 1L-21P40 and IFN-α2, both involved in extracellular matrix remodeling [85–87]. Further, the presence of neurons resulted in higher expression of FGF-2 and VEGF-A, two growth factors with potent proliferative effects (Fig. 9g).

In HGoC, the presence of neurons drastically reduced the secretion of all isoforms of the immunomodulatory transforming growth factor (TGF)-β family besides platelet-derived growth factor (PDGF)-BB (Fig. 9f). However, they persist in keeping their immunomodulatory effects by increasing the secretion of IL-1Ra, which antagonizes the inflammatory effects of IL-1 [88] and IL-13 potent immunomodulators [89, 90]. Interestingly, TGF-β1 was among a few upregulated proteins in cluster 2 of the proteomic heatmap, in which both 3D R and human GBM patients showed higher expression than 2D R, 2D NR and 3D NR (Supplementary Fig. 6a). Despite few protein numbers in this cluster, protein-protein interaction analysis constructed by the differentially expressed genes in C2 (Supplementary Fig. 6b) showed degree one interaction between TGF-β1, S100-A4, and Collagen Type I Alpha 1 Chain (COL1A1) in this cluster. Through this interaction, TGF-β1 regulates the expression and release of collagen by inducting S100 expression [91–93], where collagen forms an impenetrable barrier to protect cancerous cells from the immune system [94].

Comparing GoC with NGoC, a higher level of TGF-β in GoC explains the same radius of invasion despite different spheroid growth. In addition, in GoC, a significant increase in the level of anti-inflammatory cytokines, including Fms-like tyrosine kinase 3 ligand (Flt3L), TGF-β2, and PDGF-BB were detected, compared with NGoC (Fig. 9h). Further, a significant decrease in the level of pro-inflammatory cytokines such as IL-6, IL-8, IL-1β, and Gro-alpha in GoC were detected, compared with NGoC (Fig 9h). These findings support the decreased inflammatory reactions we observed in proteomic studies of resistant spheroids and also human recurrent GBM patient cells (Fig 7b, C4).

All in all, resistant spheroids in GoC showed higher invasiveness than non-resistant spheroids in NGoC despite their lower spheroid growth. In addition, they showed lower inflammatory reactions similar to HR. The presence of neurons resulted in a higher spheroid growth of non-resistant spheroids in NHGoC, but lower invasiveness of resistant spheroids in HGoC.

## Discussion

In this work, we applied multi-omics approach to a microfluidic 3D model of TMZ-resistant glioblastoma to gain insight into cellular pathways involved in tumor resistance. Initially, we confirmed that the 3D model recapitulated some aspects of the pathophysiology of human recurrent glioblastoma including amoeboid phenotype, mitotic quiescence, OXPHOS metabolism, and lower apoptotic cell death. Importantly, we showcased the capability of our model to understand the mechanisms underlying chemoresistance. Additionally, we showed that our model resembles some hallmarks of recurrent human glioblastoma by mirroring molecular signatures observed in recurrent GBM patient tissues, including altered cholesterol pathway and increased vesicular transport [74].

Here, we showed EMT, mitotic quiescence, OXPHOS metabolism, and inhibition of autophagy flux is associated with acquired resistance. EMT is a characteristic of cancer stem cells, more specifically mesenchymal stem cells, and has been introduced as a mechanism of chemoresistance for more than a decade in different types of cancers [60, 95] and more specifically in glioblastoma [96]. This phenotype was accompanied by mitotic quiescence, OXPHOS metabolism, and autophagy flux inhibition. Similarly, an *in vitro* study reported the reversion of EMT by autophagy induction [97], and a recent review introduced autophagy as the modulator of the EMT in glioma cells [65]. In addition, the individual role of autophagy on glioblastoma chemoresistance is reported frequently [14, 64, 98]. Mitotic quiescence is frequently reported in chemotherapy-resistant cancer stem cells [99, 100]. In line with our finding, Vlashi and colleagues were the first group reporting the rely of resistant glioma stem cells on oxidative phosphorylation [67]. Later, Viale and colleagues showed a subpopulation of pancreatic cancer cells with dormant features dependent on mitochondrial respiration responsible for tumor relapse too [101]. A recent discovery on the role of a mitochondrial receptor on the glioma stem cell resistance via oxidative phosphorylation further highlights the importance of OXPHOS metabolism on glioblastoma resistance [102]. Interestingly, Liang and colleagues introduced inhibition of the lysosomal activity, the predisposing mechanism that preserves stem cell dormancy [103].

We evaluated the effect of culturing glioma cells in 3D on their protein profile. Our results suggest that glioma spheroids exhibited higher levels of mitochondrial oxidation and cell viability with lower secretion of proinflamatory cytokines and attachment properties that resulted in MAT. Uncovering the critical role of culture conditions, our study revealed enhanced mitochondrial oxidation, mainly OXPHOS in resistant spheroids and fatty acid beta oxidation in non-resistant spheroids, a significant finding in cells cultured under 3D conditions. OXPHOS provides higher level of energy for cells, and unsurprisingly 3D culture was associated with drug-resistance in cancer spheroids through higher oxidative drug extrusion [104]. A recent proteomic study comparing drug-resistant melanoma cells and spheroids displayed higher OXPHOS in spheroids [105]. Comparable results are reported for colorectal cancer and pancreatic ductal adenocarcinoma [106]. Mesenchymal-amoeboid transition in resistant cells cultured in 3D is our another finding. MAT is introduced as a cellular state in EMT program to help mesenchymal cell increase their migration velocity [107]. Higher invasiveness of amoeboid state is attributed to higher secretion of several matrix metalloproteinases including collagenase and gelatinase [108]. In parallel with our results, Liu and colleagues applied confined condition with low focal adhesion to convert mesenchymal cells to fast migratory amoeboid cells [73]. Chikina and colleagues studying the mesenchymal fibrosarcoma cells displayed transition to amoeboid state in a population with stem cell feature [109]. Ek and colleagues, using a 3D model of hepatocellular carcinoma (HCC) and colorectal carcinoma (CRC) reported higher sensitivity of the 3D model cells to OXPHOS inhibitor compared to the monolayer cells while 3D models were more resistant to chemotherapeutic agents [110].

To further validate our model, we showed that TMZ-resistant spheroids grown in 3D model mirrored some molecular signatures of human recurrent glioblastoma. Specifically, the cholesterol pathway was significantly affected, vesicular transport was increased, and mitosis was quiescence. Further, we showed that increased vesicular trafficking at least in part is due to inhibition of autophagy flux. Studying the paired and recurrent glioblastoma patient tissues, Dekker and colleagues introduced the cholesterol metabolism and synaptic vesicular pathways as the few pathways that were affected at the different omics levels in recurrent human glioblastoma [74]. Cholesterol plays a critical role in vesicular trafficking [111] by affecting membrane fusion [112]. A whole-cell membrane capacitance measurement showed that extraction of cholesterol caused impaired vesicular endocytosis [113] and exocytosis by controlling the fusion pore conductance [114]. On the other hand, cellular cholesterol level is regulated by endocytosis of cellular cholesterol content to lysosomes [115], where cholesterol induces the activation of mTOR [116, 117], a known regulator of cell translation and proliferation [118, 119]. The similar mutual cross-talk and regulatory effect between cholesterol and autophagy pathways has been defined. Cholesterol induces autophagy [120–122], and autophagy plays an important role on cholesterol homeostasis via cholesterol trafficking [123, 124]. Thus, targeting cholesterol pathway has been introduced as a new therapeutic approach to combat glioblastoma [125–127]. In addition, the mitotic quiescence of resistant cancer stem cells has been reported frequently in different type of cancers from prostate cancer [128] and breast cancer [99] to brain tumors [129] and many other types [100, 130]. Similarly, many studies have reported the role of the autophagy pathway on the development of chemoresistance [63, 131–133].

Finally, we have exhibited how HGoC simulates the microphysiology of the glioblastoma tumor microenvironment by directly co-culturing healthy human neurons with resistant glioblastoma spheroids that resulted in higher growth of resistant spheroids while declined their invasiveness. In a same way, Venkatesh and colleagues have shown that neuronal activity augments glioma growth by induction of PI3K-mTOR pathway via secretion of neuroglin-3, a synaptic protein [134]. This group later presented data on failure of patient-derived pediatric GBM cell growth in an orthotopic xenografts mice model of pediatric GBM with neuroglin-3 knockout [135]. To our knowledge, we are the first group who has shown the effect of glioma-neuronal interaction on glioma cell invasiveness. Tang and colleagues applied neural stem cells, macrophages, astrocytes and glioma stem cells in a 3D bioprinted model and have shown the effect of the presence of macrophages, but not neurons, on glioma invasiveness [136]. Fuchs and colleagues co-cultured pediatric glioblastoma with glutamatergic neurons in a compartmentalized microfluidic chip to evaluate their electrical interaction [137], but not the effect of this interaction on glioma cell growth and invasion. Further, they cultured neurons in a separate channel that hinders direct interaction between these two types of the cells.

Despite all observed similarities, our model limited to using neuronal cells to recapitulate tumor surrounding microenvironment. However, the glioblastoma tumor microenvironment consists of other cells, including tumor associated macrophages (mostly microglia, infiltrated macrophages and monocytes) that affect tumor by both direct cell-cell interaction and release of cytokines and chemokines and matrix metalloproteinases [138]. Through these interactions macrophages also impact on brain tumor’s growth and invasion [139]. In addition to macrophages, other innate immune cells including neutrophils and mast cells and adaptive immune cells including lymphocytes, are recruited to the tumor microenvironment too [138]. Reciprocal interaction between the immune cells and GBM cells play a prominent role on the glioma behaviour [138]. Astrocytes are other cells recruited to the GBM microenvironment and significantly modify the GBM growth [140], via both cell-cell interaction and extracellular vesicles [141]. The blood-brain barrier is another important factor that affects the treatment of brain tumors due to restricting the passage of molecules from vessels to the brain and tumor subsequently, and pericytes has been introduced as the major agent affecting chemoresistance in GBM besides to the endothelial cells [142]. The other limitation of our study is using a cell line that inherently results in differences in some aspects with human samples. Our final goal is applying human recurrent samples to develop spheroids and incorporating other cells including neurons, microglia and astrocytes in direct co-culture with spheroids in the microenvironment channel. Additionally, we will culture endothelial cells and pericytes in the blood channel to mimic the blood-brain barrier, and lymphocytes to nutrition media to better mimic the human blood vesicles.

Taken together, our study exhibited that our hybrid HGoC model overcomes limitations in spheroid and microfluidic models and simulates critical features of micropathophysiology of human recurrent glioblastoma, including altered cholesterol metabolism and increased vesicular trafficking, that our data suggest it might be due to an inhibited autophagy flux. HGoC enables a systematic evaluation of mechanism of TMZ-resistance and disclosed mitotic quiescence, OXPHOS metabolism, and MAT, characters of cancer stem cell, as part of mechanism of TMZ-resistance that can be targeted as a novel approach to overcome GBM recurrence. Thus, HGoC can be used to further study the mechanism of chemoresistance and discover new therapeutic approaches. In addition, it is applicable to a spectrum of other type of cancers.

## Methods

### Fabrication of Self filling Microwell array (SFMWA)

SFMWA were fabricated by replica molding technique using our 3D printed mold as described earlier [41]. In brief, 2% agarose (Molecular Biology Grade) was prepared in PBS solution and heated to 100°C to dissolve the agarose completely. The solution was then kept warm at 70 °C. 1000 µL of the solution was cast onto the microwell mold and was left at RT for 5 min. When the solution solidified, it was removed from the mold using a bent-tip spatula, washed with 70% ethanol under sterile conditions, and exposed to ultraviolet (UV) radiation for 30 minutes.

### HGoC fabrication

HGoC comprised of a TME channel 15000-μm-long, 1500-μm-width, and 400-μm-height separated from two 15000-μm-long, 3000-μm-width, and 400-μm-height blood channels with 300 µm polydimethylsiloxane (PDMS) posts. HGoC was fabricated using two-layer photolithography. Two SU-8 100 layers (MicroChem Corp) were coated on a microscope slide to achieve a thickness of 400 µm in height. One minute of corona treatment was done on each clean glass slide, followed by coating the first layer of SU-8 100 on the slide using a G3 spin coater at a speed of 1300 round per minute (RPM), followed by 30 min pre-bake at 65 °C and 90 min bake at 95 °C. The second layer of SU-8 100 was added on the top of the first layer similarly. Both layers were cross-linked in one step by 300 sec of 350 watts UV exposure while covering them with the microchannel mask. This step was followed by 1 min of pre-bake at 65 °C and 20 min at 95 °C. Development of the master mold was done by a SU-8 developer (MicroChem) for 10 min and completed by washing with isopropanol and cleaning with an air gun. The PDMS elastomer was mixed with a cross-linker (Sylgard 184 Silicone Elastomer Kit, Ellsworth Adhesives) on a ratio of 1:10. The mixture was degassed in a desiccator, poured on the SU-8 master mold, and cured for 2 hours at 80 °C. Cured PDMS was gently peeled off the mold and punched with a 1 mm and 5 mm diameter puncher to create inlets and outlets for TME and blood channels and cut into 20 × 20 mm. PDMS chips were cleaned with 100% ethanol and an air gun. Then, they were treated with oxygen plasma (Plasma cleaner 117125, Diener Zepto one) for 38 seconds, bonded to coverslips, and baked at 60 °C overnight. To be sterilized, the HGoC were soaked in 70% ethanol for 20 minutes, rinsed with absolute ethanol, dried, and exposed to UV light for 30 minutes. Finally, HGoC were baked at 80 °C for 4 hours in a sterile vented container to retain the hydrophobicity.

### Establishment of TMZ-resistant cells

U251-mKate cells (developed by Dr. Marcel Bally, Experimental Therapeutics, British Columbia Cancer Agency, Vancouver, BC, Canada) were cultured in T75 tissue culture flasks in a complete media containing high glucose DMEM (Gibco; Cat #: 11965092), 10% FBS (Gibco; Cat #: 10437-036), and 1% Penicillin-streptomycin (Gibco, Cat#:15140122) and incubated at 37°C, 7.5% CO_2_, in a humidified incubator (Thermo Fischer Scientific). We chose a pulsed-selection strategy to make a clinically relevant chemoresistant model [40]. After reaching 70-80% confluence, cells were treated with 100 µM of TMZ (Sigma-Aldrich, CAS #:85622-93-1) over three weeks, followed by four weeks of TMZ-free media for recovery. Then, cells were cultured in complete media with 250 µM TMZ for three weeks, followed by another four weeks of culture in TMZ-free media. The final population of U251-mKate cells resistant to the 250 µM TMZ was selected and cultured in a complete media without TMZ for four weeks.

### 2D culture

For all experiments, including proteomics, lipidomics, western blotting, and gene expression analysis, 10 mL of 5 × 10^5^ cell mL^−1^ of U251-mKate (NR) and TMZR-U251-mKate (R) cells were seeded in 100 mm Petri dishes and were nourished with TMZ-free and TMZ 250 μM-containing complete high glucose DMEM media, respectively. Cells were incubated at 37 °C and 7.5% CO_2_ in a humidified incubator for four days. The cells were scraped and washed using PBS. Cell pellets were kept in a PBS buffer containing a 1:75 ratio of phosphatase inhibitor cocktail (Sigma, P5726) and protease inhibitor cocktail (Sigma, P8340) at −80°C until further analysis.

### 3D culture

Spheroids were formed as described previously [41]. Briefly, sterile MWAs were transferred to the 12 well plates under sterile conditions. Then, 500 μL of 10^6^ cell mL^−1^ of R or NR cell suspension were gradually added to the loading chamber of MWAs, and the plates were transferred to the CO_2_ incubator. After one hour, the plates were removed from the incubator, and 2 mL of the complete DMEM media were gently added to the wall of each well. MWAs were incubated for four days to ensure the formation of aggregated spheroids while half of the media were changed every other day. Day 4 was selected based on our previous study showing that cells make the maximum cell junctions, confirmed by the reduction in the diameter of spheroids (Supplementary Fig. 7), over the first four days and start proliferation [143]. On day 4, cell media were aspirated and replaced with TMZ-free or TMZ 250 μM-containing complete media for 3D NR and 3D R, respectively, for four days. For GoC culture on day 4, spheroids were washed from the MWA by pipetting fresh media (details in HGoC culture session).

### Differentiation of neural progenitor cells to neurons

iPSC-derived Neural Progenitor Cells (NPC) (ATCC®, ACS-5003™) were cultured and differentiated toward neurons based on the ATCC protocol. In short, 12 well plates were coated with 0.5 mL of 150 μg mL^−1^ of CellMatrix basement membrane gel (ATCC®, ACS-3035) for one hour at 37 °C. After one hour, the coating agent was removed, and 1.5 mL of NPC growth media (DMEM-F12, (ATCC®, 30-2006)) containing NPC growth kit (ATCC®, ACS-3003) were immediately added to each well and incubated inside the CO_2_ incubator for 15 minutes. Cryopreserved NPCs were thawed in a 37 °C water bath, and cells were transferred to a 15 mL conical tube containing 9 mL DMEM-F12. Cell pellets were resuspended in NPC growth media after 5 minutes of centrifugation at 270 × g. Cells were counted and adjusted to 1.6 × 10^5^ viable cell mL^−1,^ and 0.5 mL of NPC cell suspension was added to each well of 12 well plates. The plates were returned to the incubator. Their media were gently changed 100% the next day and every other day. NPCs were passaged when they reached 95% confluence to have enough cells for differentiation. NPCs were seeded in a coated 12 well plate in a concentration of 10^5^ cell cm^−2^ and incubated overnight in NPC growth media. Then, the NPC growth media were replaced with 1.5 mL of pre-warmed differentiation media (DMEM-F12 containing Dopaminergic Neuron Differentiation Kit, ATCC® ACS-3004). Cells were incubated for three weeks for complete differentiation, while 85% of media were gently replaced with pre-warmed differentiation media every other day.

### HGoC culture

Sodium alginate was first reconstituted at 4% in a medium with 150 mM NaCl. This solution was then incubated for six hours at 37 °C to enable further dissolution of the alginate. The alginate stock solution was passed through a 0.22 μm filter and kept on ice. The CaCl_2_ solution was individually passed through 0.22 μm filters and kept on ice. An aliquot of Matrigel was kept on ice to be thawed. Spheroids were separated from the MWA by washing using the media. The spheroids were centrifuged and resuspended in complete cold DMEM-F12. A pipette tip was chilled by pipetting cold serum-free DMEM. This pipette tip was used to resuspend 10^5^ neurons in 21.5 μL of spheroids in complete DMEM-F12. The spheroid-neuron combination was mixed with 65 μL of Matrigel in a 1.5 mL centrifuge tube using a chilled pipette tip. Then 12.5μL of alginate 4% was mixed with the mixture. Finally, 1.2 μL of CaCl_2_ 1 M was added to make a cell-spheroid combination encapsulated in 4.6 mg mL^−1^ of Matrigel and 5 mg mL^−1^ of alginate with 12.5 mM of CaCl_2_ as the cross-linker. The mixture was vortexed for 10 s and kept on ice during the experiment. 10 μL of the mixture were injected into the TME channel of each microfluidic chip. All steps were done using chilled pipette tips. Immediately after injection, chips were filliped upside down to avoid attachment of spheroids to the bottom of the TME channel and were transferred to the CO_2_ incubator. After 45 minutes, they were removed from the incubator and flapped. Then, complete differentiation media were added to the blood channels, and the HGoC chips were returned to the incubator for later experiments.

### Viability assay

According to the manufacturer’s protocol, a viability assay was performed using PrestoBlue (Invitrogen, A13262). In brief, 100 μL of 5 × 10^5^ cell mL^−1^ of R or NR cells were cultured in 96 well plates and treated with TMZ-free or TMZ 250 μM-containing complete high glucose DMEM media for one day. Then the media was removed, and cells were treated with 0-5000 μM of TMZ for 96 hrs. A working solution of PrestoBlue was prepared with a 1:10 dilution of stock solution. After four days of treatment, the media was replaced with 200 µL of working solution, and the plates were incubated in the CO_2_ incubator for 30 min. For 3D culture, on day 4 of the formation of the spheroids on MWA, the media was replaced with 1 mL of working solution in the 12 well plates, and the plates were incubated in the CO_2_ incubator overnight. Once incubated, 150 µL of each sample was transferred to a 96 well plate. The fluorescent intensity was read using an Infinite M Nano plate reader (Tecan), via the i-control software (Tecan, Version 3.9.1.0), at two wavelengths: 560 nm and 590 nm.

### Live/dead imaging

For live/dead imaging, confocal fluorescence microscopy was used for samples stained with the L/D kit (Invitrogen, MP 03224). The L/D staining solution, 1 μM calcein AM and 4 μM ethidium homodimer-1, were prepared according to the supplier’s protocol. For 2D culture, cells were cultured in 12 well plates until they reached 80% confluence. Then, the media were removed, and 200 μL of L/D solution was added to the wells immediately. The plates were incubated at RT in the dark for two hours, and the confocal imaging was done immediately. For L/D imaging of 3D R and 3D NR spheroids, on day four of formation, they were washed from MWA, and 2-3 spheroids were transferred to a well of 96 well plates. 100 μL of L/D solution was added to the wells, and the plates were incubated at RT in the dark for two hours. The confocal imaging was done once incubation time was finished. For L/D imaging of HGoC, the media were removed from the blood channel, and L/D solution was added to the blood channel. The HGoC was incubated at RT in the dark for two hours, and the confocal imaging was performed immediately.

### Immunofluorescence Imaging

Cells were fixed with 3.7% formaldehyde (VWR) and incubated at RT for 30 min. The samples were washed three times using DPBS, while kept at DPBS for 5 min each time. Fixed samples were permeabilized and blocked overnight at 4 °C with a blocking buffer containing 5% IgG-free bovine serum albumin (BSA) (Jackson ImmunoResearch) and 0.3% Triton X-100 (Bio Basic) in DPBS. The blocking buffer was replaced with an antibody solution containing 1% BSA, 0.3% Triton X-100 and conjugated antibodies in DPBS and incubated overnight at 4 °C in the dark. The dilution ratio of 1:200 for Cleaved PARP (Cell Signaling Technology, Asp214, D64E10), 1:100 for Vimentin (Cell Signaling Technology, D21H3), and 1:2000 for LC3 (Sigma, L7543) were used. For LC3, that primary antibody was not conjugated. After removing the primary antibody, a secondary antibody at 1:2000 dilution (Jackson ImmunoResearch Laboratories, 711-545-152) was applied and incubated at RT for 60 min in the dark. For actin cytoskeleton (Thermo Fischer Scientific, A12379) a 1:1000 dilution was used. Then, the samples were washed three times with DPBS, and the solution of To-pro3 (Thermo Fischer Scientific, T3605, 1:1000) or Hoechst (Biotium, 33342, 5μM) was added and incubated for 30 min at RT in the dark to stain the nuclei. Finally, the nuclei staining solution was removed, and the samples were washed with DPBS three times, incubating for 5 min. The wells were partially filled with DBPS, and the samples were kept at 4 °C in the dark until imaging was performed. All staining solutions were introduced to HGoC through the blood channels, and 2D and 3D were added to the well plates.

### MitoView Live Imaging

MitoView Green (Biotium, 70054) and Hoechst (Biotium, 33342) were used at 100 nM and 5 µM to identify mitochondrial morphology and nuclei, respectively. Cells were stained and incubated for 15-30 min at 37 °C.

### Imaging

The bright-field (BF) images were captured by Axio Observer ZEISS microscope (ZEISS, Oberkochen, Germany). For LC3 and Mitoview, Zeiss Axiovert 200 inverted microscope fitted with a Calibri 7 LED Light Source and Axiocam 702 mono camera were used. Quantification, scale bars, background subtraction, and processing were done on Fiji (ImageJ) and Zen 2.3 Pro software. HGoC imaging was performed using a ZEISS confocal microscope (Zeiss LSM880, ZEISS, Oberkochen, Germany) with 20x and 50x magnification objectives.

### Gene expression assay

Based on the manufacturer protocol, RNA extraction was performed using the PureLink RNA Mini kit (Invitrogen, 12183018A). 500 ng of total RNA was used to synthesize cDNA using the LunaScript RT SuperMix Kit (New England Biolabs, E3010L). The Luna Universal qPCR Master Mix (New England Biolabs, M3003X) was used to monitor cDNA amplification, 10 ng/well in 96 well plate format, on a real-time PCR machine (Bio-Rad, CFX96). To evaluate the expression of LDL-R and SREBP-2 genes, LDL-R forward primer, 5’-CCCGACCCCTACCCACTT-3’; reverse primer, 5’-AATAACACAAATGCCAAATGTACACA-3’; and SREBP-2 forward primer, 5’-GACGCCAAGATGCACAAGTC-3’; reverse primer, 5’-ACCAGACTGCCTAGGTCGAT-3’ were used. The expression of β-ACTIN forward primer, 5’-CACCATTGGCAATGAGCGGTT-3’; reverse primer, 5’-AGGTCTTTGCGGATGTCCACGT-3’; EIF2a forward primer, 5’-CGAAACACTGTCTCTCAGTCAA-3’; reverse primer, 5’-CCAGTTGCTGCTTGTTCTTTC-3’; and 18S forward primer, 5’-GCAGAATCCACGCCAGTACAAG-3’; reverse primer 5’-GCTTGTTGTCCAGACCATTGGC-3’ were evaluated as the reference genes. A geomean of β-ACTIN, EIF2a, 18S was used to normalize gene expression. Primer specificity was tested in silico PCR using ENCODE UCSC database, and PCR products on 1% agarose gel made with Tris-Acetate-EDTA buffer diluted to 1X (TAW50X01, MP Biomedicals) at 100V for 45 min.

### Western blotting

Immunoblotting was performed as described previously [144]. Antibodies were used as follows: primary anti-LC3 (Cell Signaling Technology, 2775S), primary anti-p62 (Cell Signaling Technology, 5114S), and secondary anti-body was anti-Rabbit (Sigma, A6154). Ponceau S stain was used to validate equal loading.

### Cholesterol mass measurement

2D R and 2D NR cells were cultured in cholesterol-free media and harvested after 24, 48, and 72 hours. According to the manufacturer’s instructions, cellular cholesterol content was determined as described previously [144, 145]using the Amplex Red CH assay kit (Invitrogen, Thermofisher, Toronto, Canada).

### Cholesterol ester assay

De Novo Cholesterol biosynthesis was measured using a modified Mokashi protocol [145]. In brief, 2D R and 2D NR cells were cultured in cholesterol-free media containing ITS (insulin transferrin selenium) for 24, 48, and 72 hours. Then, cells were treated with serum-free DMEM containing 2 µCi [^14^C] acetate per 2-mL dish for the indicated times; then, the cells were harvested and washed twice with ice-cold PBS (pH 7.4). Total lipids were extracted [146], and the incorporation of radioisotopes into cholesterol and cholesteryl ester were determined and normalized to protein content as previously described [147].

### Transmission electron microscopy

To show the structural changes associated with resistant development, TEM was performed using a protocol described in detail previously [148, 149]. Briefly, 2D R and 2D NR cells were cultured in 100 mm dishes and treated per OEP for 72 h. Ultra-thin sections (100 nm on 200 mesh grids) were used 36 hours after treatment for uranyl acetate staining, and were then counterstained with lead citrate.

### Invasion assay

Invasion of the spheroids to the surrounding tissue is imaged by an inverted microscope’s 10x and 20x objective lenses (Supplementary Fig. 8a). Spheroids’ diameter and invasion length were measured using ImageJ [150]. The Freehand Selection Tool was used to trace the boundary of the tumor to exclude the invasion area from the core of the spheroid. A circle was then fitted on the marked boundary to approximate the spheroid’s core to a perfect sphere. The diameter of the marked circle was measured by the software’s Feret’s Diameter measurement tool (Supplementary Fig. 8b). To identify the invasions, the active image was converted to 32-bit grayscale. This conversion enables the software to segment the grayscale images into features of interest by interactively setting upper and lower values. After the segmentation of invasions, the Wand Tracing tool was used to highlight the boundary of the invasions (Supplementary Fig. 8b).

### Measurement of mitochondrial respiration

Mitochondrial respiration was measured using the Agilent Seahorse XFe24 analyzer by recording the Oxygen consumption rate (OCR). Approximately 0.3 × 10^6^ cells (NR and R) per well were seeded in their growth medium in a XF cell culture 24 well microplates. On the day of the experiment, cells were washed twice with and changed to XF base minimal DMEM media containing glucose, L-glutamine and sodium pyruvate (Agilent tech #103334-100). The cells were incubated at 37 °C for 1 hr prior to starting the assay. Seahorse analyzer measured the baseline OCR first, followed by proton leak using 1 µM of oligomycin. Maximal respiration was determined by injection of FCCP (2 µM) and non-mitochondrial respiration by addition of 1 µM rotenone and 1 µM antimycin A together. OCR date was normalized by live cell count signal obtained by staining the individual wells with Hoechst33342 (Invitrogen #R37605) using Cytation 5 (Biotek). Maximal respiration was calculated by subtracting non-mitochondrial OCR from FCCP OCR. Spare capacity was determined by subtracting baseline OCR from FCCP OCR. Further, ATP production was calculated by subtracting Oligomycin OCR from baseline OCR. Finally, proton leak was measured by subtracting non-mitochondrial OCR from Oligomycin OCR. Data are presented as pmol of oxygen/minute/cell count.

### Liquid chromatography tandem mass spectrometry analysis of peptide mixture

An aliquot of each sample was removed, and a Bradford assay was used to determine protein concentration. A volume from each sample containing 15 µg of protein was pipetted into a solubilizing buffer of 100 mM Tris/1% sodium deoxycholate and incubated at 95 °C for 10 minutes. The protein was then digested with Lys-C/Trypsin and purified using a Preomics (Martinsried, DE) iST sample preparation kit.

The PreOmics kit digested samples were separated by two-column online reverse phase chromatography using a Thermo Scientific EASY-nLC 1000 system with a reversed-phase pre-column Magic C18-AQ (100 µm ID, 2.5 cm length, 5 µm, 100 Å) and an in-house prepared reversed-phase nano-analytical column Magic C-18AQ (75 µm ID, 15 cm length, 5 µm, 100 Å, Michrom BioResources Inc, Auburn, CA) at a flow rate of 300 nl min^−1^. The chromatography system was coupled online with an Orbitrap Fusion Tribrid mass spectrometer (Thermo Fisher Scientific, San Jose, CA) and a Nanospray Flex NG source (Thermo Fisher Scientific). Solvents were A: 0.1% Formic acid/Water; B: 90% Acetonitrile, 10% Water, 0.1% Formic acid. After a 348 bar (~ 3µL) pre-column equilibration with a 348 bar (~ 4µL) nanocolumn equilibration, 2 µL of the sample was injected and pressure loaded at 300 bar for a 6 µL desalting. The peptides were then separated by a 126 min analysis (0 min: 5% B; 100 min: 24% B; 120 min: 38% B; 121 min: 100% B; 126 min: 100% B).

The Orbitrap Fusion 3.1 instrument parameters were as follows for the Orbitrap (OT-MS) with Ion-trap MS2 (IT-HCD) analysis: Nano-electrospray ion source with spray voltage 2.55 kV, capillary temperature 275 ℃. Survey MS1 scan m/z range 300-1500 profile mode, resolution 120,000 FWHM@200 m/z one Microscan with maximum inject time 50 ms. The Siloxane mass 445.120024 was used as lock mass for internal calibration. Data-dependent acquisition Orbitrap survey spectra were scheduled at least every 3 seconds, with the software determining the “Top-speed” number of MS/MS acquisitions during this period. The automatic gain control (AGC) target values for FTMS and MSn were 400,000 and 10,000 (MS2), respectively. The most intense ions charge state 2-7 exceeding 10,000 counts were selected for HCD MSMS fragmentation with orbitrap detection in centroid mode. Monoisotopic Precursor Selection (MIPS) was enabled with Dynamic exclusion settings: repeat count: 1; exclusion duration: 15 seconds with a 10 ppm mass window. The ddMS2 IT HCD scan used a quadrupole isolation window of 1.6 Da; auto-scan range with rapid scan rate, centroid detection, first mass 110 m/z, 1 Microscan, 246 ms maximum injection time, and a stepped HCD collision energy setting of 28,30, and 32% [151].

### Proteomics Data Analysis

The mass spectrometry proteomics data have been deposited to the ProteomeXchange Consortium via the PRIDE [152] partner repository with the dataset identifier PXD037733. Raw files were created using the XCalibur 4.2.28.14 (Thermo Scientific) software. Thermo Raw files were uploaded to MaxQuant (V.1.6.2.10), and MS/MS spectra were searched against the Uniprot (Swiss-Prot) protein sequence database (Homo Sapiens, release version of May 2021) using Andromeda, with an FDR for identification set at 0.1. The MaxLFQ algorithm quantified proteins. Database search parameters were as follows: protein N-term acetylation and deamidation, methionine oxidation as variable modifications, and cysteine carbamidomethylation as fixed modification. As a result, 4122 identified proteins were uploaded into Perseus (V.1.6.2.2). Data were filtered to remove proteins identified in the reverse sequences with only modified peptides considered common lab contaminants and annotated, resulting in 3879 proteins. For the Principal Component Analysis (PCA) and Hierarchical Clustering Analysis (HCA), proteins with the non-statistical difference between sample groups, unrelated to TMZ resistance, were removed following an ANOVA analysis. The expression values of the 1100 identified proteins were normalized using Z-score. In PCA, the normalized values were plotted, considering a Benjamini-Hochberg threshold of 0.05, using the built-in tool of Perseus. In parallel, according to the Euclidean distance, HCA was performed to evaluate the relative similarities between the different groups.

The data were filtered, considering a minimum of 70% valid values in each group. The non-normalized expression values of 2024 identified proteins were log2 transformed, and missing values were replaced with the value of −1. Differentially expressed proteins between sample categories were then determined using a 2-sided t-test with 250 randomizations. The FDR was set to 0.05 and the S0 to 0.1. The data was then plotted as volcano plots. Data were exported and uploaded to JMP 16.0.0 for visualization without further processing. Upregulated proteins are shown in red, whereas downregulated are shown in blue.

Proteomics data were retrieved from the ProteomeXchange Consortium/PRIDE repository (PXD017952) for comparative analysis with human primary and recurrent tumors and analyzed in MaxQuant (V.1.6.8.0), considering an FDR set at 1% on the PSM, peptide and protein level. Spectra were compared to the human protein sequences of Uniprot (Homo Sapiens, release version of May 2021) and proteins quantified by the MaxLFQ algorithm with a minimum ratio count of two unique or razor peptides. Database search parameters were as described above. For MaxQuant, the mass tolerance of precursor mass was 20 ppm, while fragment ion mass tolerance was 0.15 Da. The minimal peptide length was defined as 7 amino acids; at most, two missed cleavages were allowed. A single matrix containing the expression of common proteins in TMZR and TMZS cells and primary and recurrent tumors (1191 proteins) was built and uploaded in Perseus (V.1.6.2.2). As mentioned, potential contaminants, reverse identification, and proteins only identified by site were removed. Samples were grouped and filtered to contain a 70% minimum of valid values per group (1007 proteins). Non-normalized expression values were log2 transformed and missing values imputed according to a normal distribution (width 0.3, downshift 1.8). Log2 values were then *Z*-scored and hierarchically clustered as aforementioned. For visualization purposes, the normalized data were exported and uploaded to MetaboAnalyst 5.0 without further processing. The heatmap was then manually curated to remove clusters that do not consider proteins involved in gaining resistance to TMZ, that is, proteins with a non-significant difference between groups.

Protein-protein interaction networks were constructed using Network Analyst [153, 154]. In brief, minimum first-order interaction networks were constructed from proteins with p<0.05 using the IMEx interactome [155]. Pathway analysis was performed on upregulated and down-regulated proteins within each network using REACTOME and Gene Ontology enrichment analysis built-in Network Analyst. Gene ontology enrichment analysis was also done on the two largest nodes with altered abundance in each network (excluding first-order interactors). Proteins with increased abundance are shown in red, and decreased abundance is shown in blue.

### Liquid chromatography tandem mass spectrometry analysis of lipid species

Lipids extraction was done using chloroform: methanol (C:M), 2:1 (v/v). Briefly, a mixture of C:M, 100 µl of cell homogenate, and 30 µL of the internal standard mixture was made. Samples were vortexed and centrifuged (3500 rpm at RT for 5 minutes). The lower phase was transferred to a different tube and dried under a stream of N_2_ gas at room temperature. Lipids were then reconstituted in 50 µL water-saturated butanol and sonicated for 10 minutes. In the last step, 50 µL of methanol with 10 mM ammonium formate was added, and samples were centrifuged (10000 g, 10 min, RT). Lipid internal standards (ISTD) were either odd-chain or deuterated and were not present endogenously. Samples were transferred into the micro insert in sample vials for lipid analysis (for details on lipid abbreviations, chemical information, and LC-MS conditions for each lipid species, refer to Supplementary Table.4).

Lipids were separated on a reverse-phase liquid chromatography-electrospray ionization tandem mass spectrometry (LC/ESI-MS/MS) platform using a Prominence chromatographic system (Shimadzu Corporation, OR, USA) as previously described [156]. Instrument control and data processing were done with Analyst v1.6 and MultiQuant v2.1 software (AB Sciex, MA, USA). The separation was achieved on a Zorbax C18, 1.8 μm, 50 × 2.1 mm column (Agilent Technologies, Mississauga, ON). The flow rate was set to 300 µL/min using a linear gradient of mobile phase A and mobile phase B. Mobile phase A and B consisted of tetrahydrofuran, methanol, and water in the ratio of 20:20:60 (v/v/v) and 75:20:5 (v/v/v), respectively. Both A and B contained 10 mM ammonium formate. The elution program was as follows: start with 0% solvent B; increase to 100% B at 8.00 min; maintained at 100% B for 2.5 min; and back to 0% B over 0.5 min. The column was finally re-equilibrated to the starting conditions (0% mobile phase B) for 3 min before the next sample injection. Diacylglycerol (DG) and triacylglycerol (TG) species were separated in an isocratic fashion (100 µL/min) by employing 85% mobile phase B over 8 min. The analytical column and autosampler were maintained during the analysis at 50°C and 25°C, respectively. The injection volume was 5 µL.

Mass Spectral Analysis: Lipids eluted from the HPLC system were introduced into the AbSciex 4000 QTRAP triple quadrupole linear hybrid mass spectrometer. The mass spectrometer was operated in scheduled Multiple Reaction Monitoring (MRM) mode. 322 unique lipids spanning 25 different lipid classes/subclasses were screened for targeted semi-quantitation. All lipid species other than fatty acids were scanned in positive electrospray ionization mode [ESI+]. The individual lipids in each lipid class were identified by lipid class-specific precursor ions or neutral losses. Lipids were then quantified by comparing the deisotoped lipid peak areas against those of the class-specific ISTDs added before lipid extraction. Lipids were represented by the total carbon number of the fatty acids. In the ESI+ mode, the instrument settings were optimised as follows: curtain gas (psi), 26; collision gas (nitrogen), medium; ion spray voltage (V), 5500; temperature (°C), 500.0; ion source gas 1 (psi), 40.0; ion and source gas 2 (psi), 30.0. The MRM detection window was fixed between 45s and 90s depending upon the chromatographic peak width of the lipid class. Isobaric species within the same class, such as PC(O) and PC(P), exhibited clear separation in this method. Also, molecular species within the same lipid class, which differ only in the number of double bonds, were well separated chromatographically.

### Lipidomics and secretome data Analysis

The samples were investigated for quartile normalization to have similar samples for data preparation before analysis. If Box plots confirmed a lack of similarities, quantile normalization was performed, and samples became similar in pattern (Supplementary Fig. 9 a & b). Since lipids had different measurement scale units, we standardized them to omit measurement scales. The pairwise comparison analysis was performed by limma package in Bioconductor to mine statistically significant lipids/cytokines. The significant lipid/cytokines were extracted based on the difference in their expression values with log_2_ fold change ≥ 2 and an adjusted p-value threshold of 0.001 [157]. All analyses were performed with R software (version 4.1.1). Limma, pheatmap, and ggplot2 packages were used.

### Statistical analysis

All experiments were repeated at least three times independently, and the values were calculated by averaging the results of the replicates. Error bars show ± standard deviation from the average values. Significance analysis was performed using a two-tailed t-test or one-way or two-way ANOVA using GraphPad Prism, followed by Bonferroni correction with p-Value=0.05.

## Data availability

The main results supporting this study are accessible within the paper and Supplementary Information. The other raw and analyzed data will be available from the corresponding authors upon request. Proteomics data are available at the ProteomeXchange Consortium/PRIDE repository via the dataset identifier PXD037733.

## Acknowledgments

This work was supported by Research Manitoba New Investigator Operating Grant and University Collaborative Research Program, BC Cancer Foundation, Canadian Foundation for Innovation, and NSERC. We acknowledge Dr. Marcel Bally, who generously gave us the U251-mKate cells, and Dr. Carla Vitorino for insights on proteomic study. We appreciate Ehsan Samiei and Erik Pagan for their contribution on designing the chip and drawing schematic respectively. João Basso acknowledges the Portuguese Foundation for Science and Technology (FCT) for the PhD research grant (SFRH/BD/149138/2019).

## Author contributions

SS, MA and SG conceptualized the study. SS, MAB, JA, CC, SDSR, MT, LC, MH performed the experiments. SS, JB, RV, CP and TD did the bioinformatics analysis. MA, SG, SS, RV, DS, AR, GH, and VD supervised the experiments. MA, SG, SS provided overall guidance and supervised the project. SS led the project and wrote the manuscript. MA, SG edited and finalized the manuscript. All authors reviewed and commented on the manuscript.

