## Supplementary Figures. for "A multi-omics analysis of glioma chemoresistance using a hybrid microphysiological model of glioblastoma": Supplementary Figures.pptx

### Slide 1
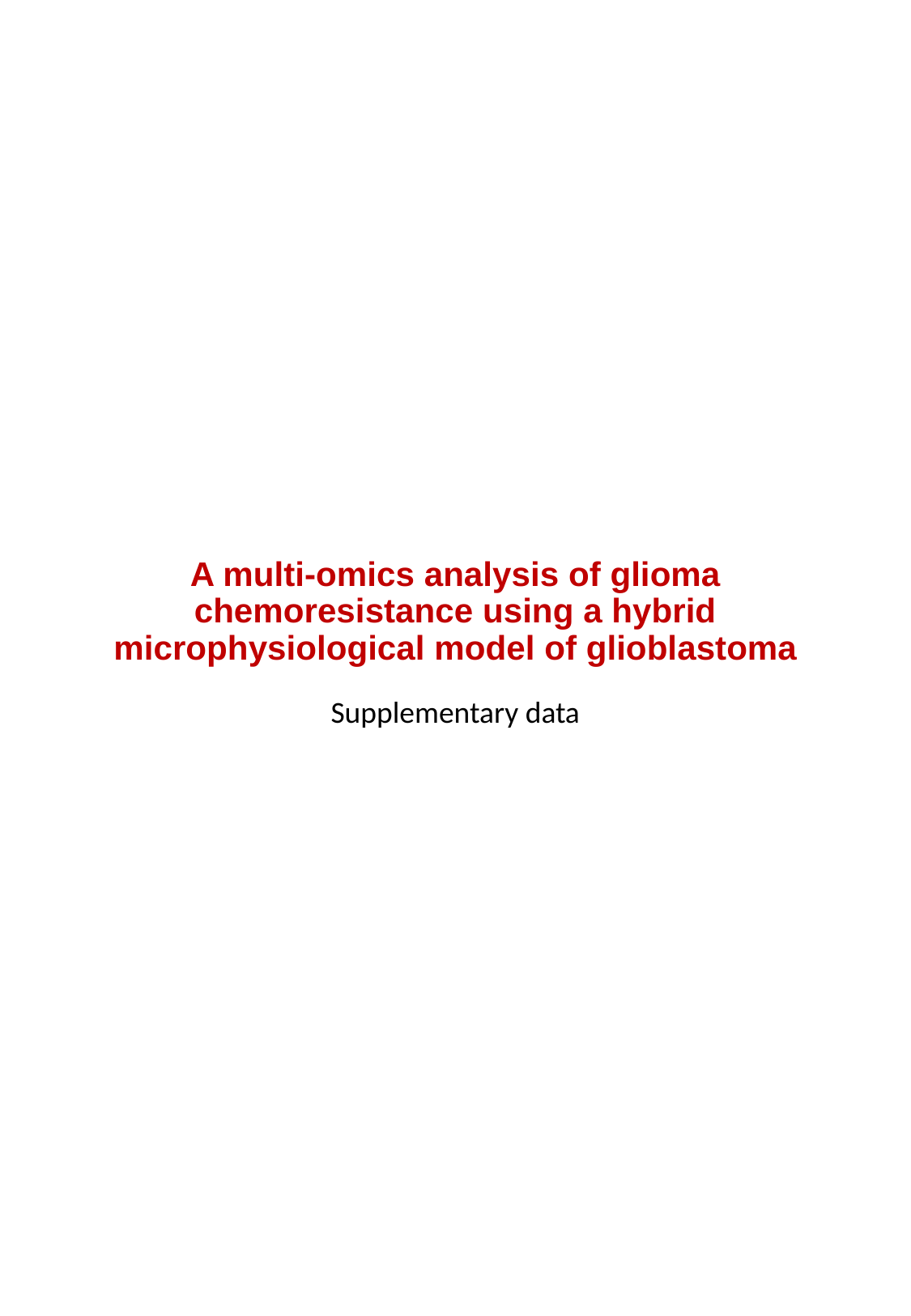

#### A multi-omics analysis of glioma chemoresistance using a hybrid microphysiological model of glioblastoma
Supplementary data

### Slide 2
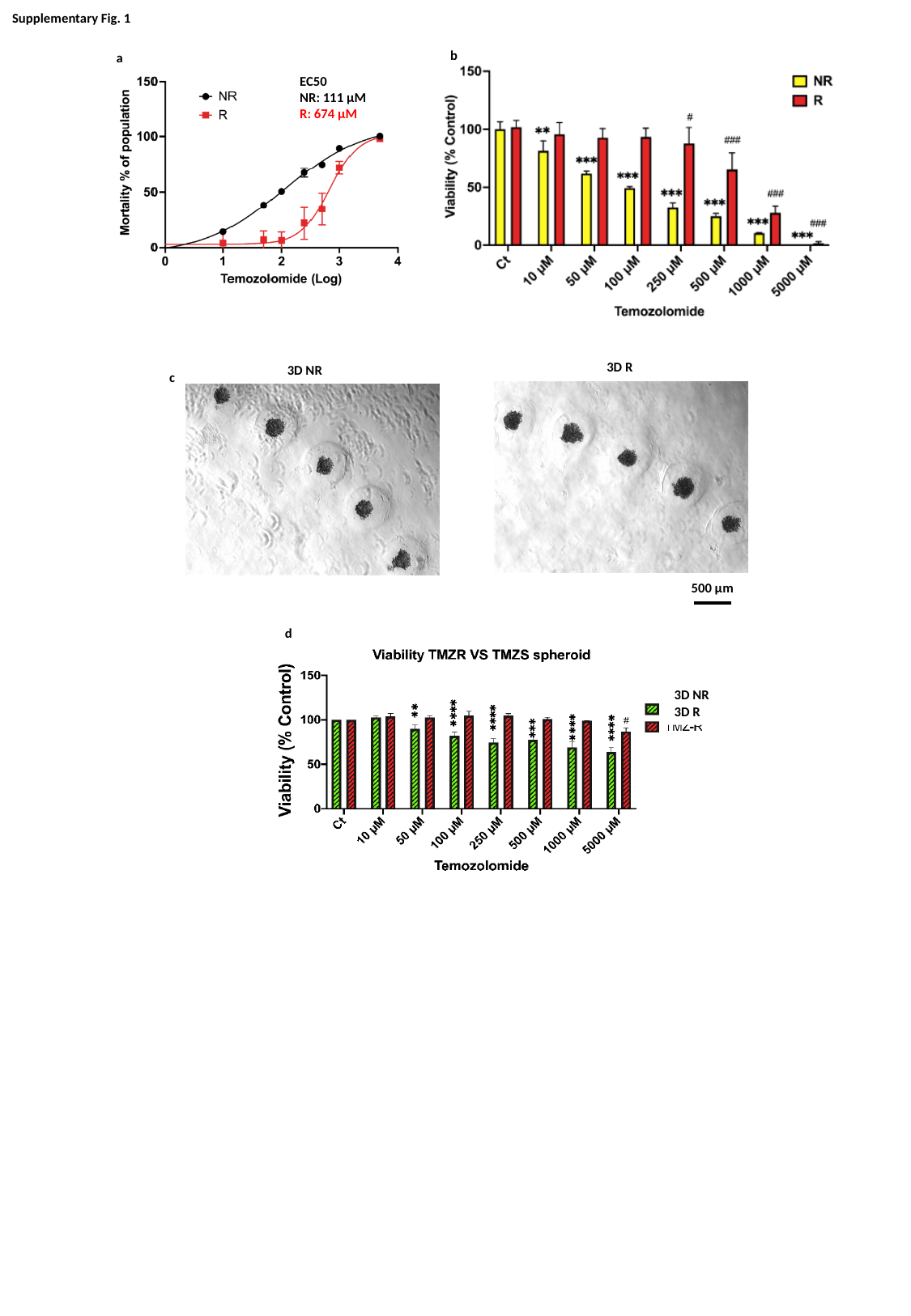

#### Supplementary Fig. 1
b
a
EC50
NR: 111 µM
R: 674 µM
3D R
3D NR
c
500 μm
d
3D NR
3D R

### Slide 3
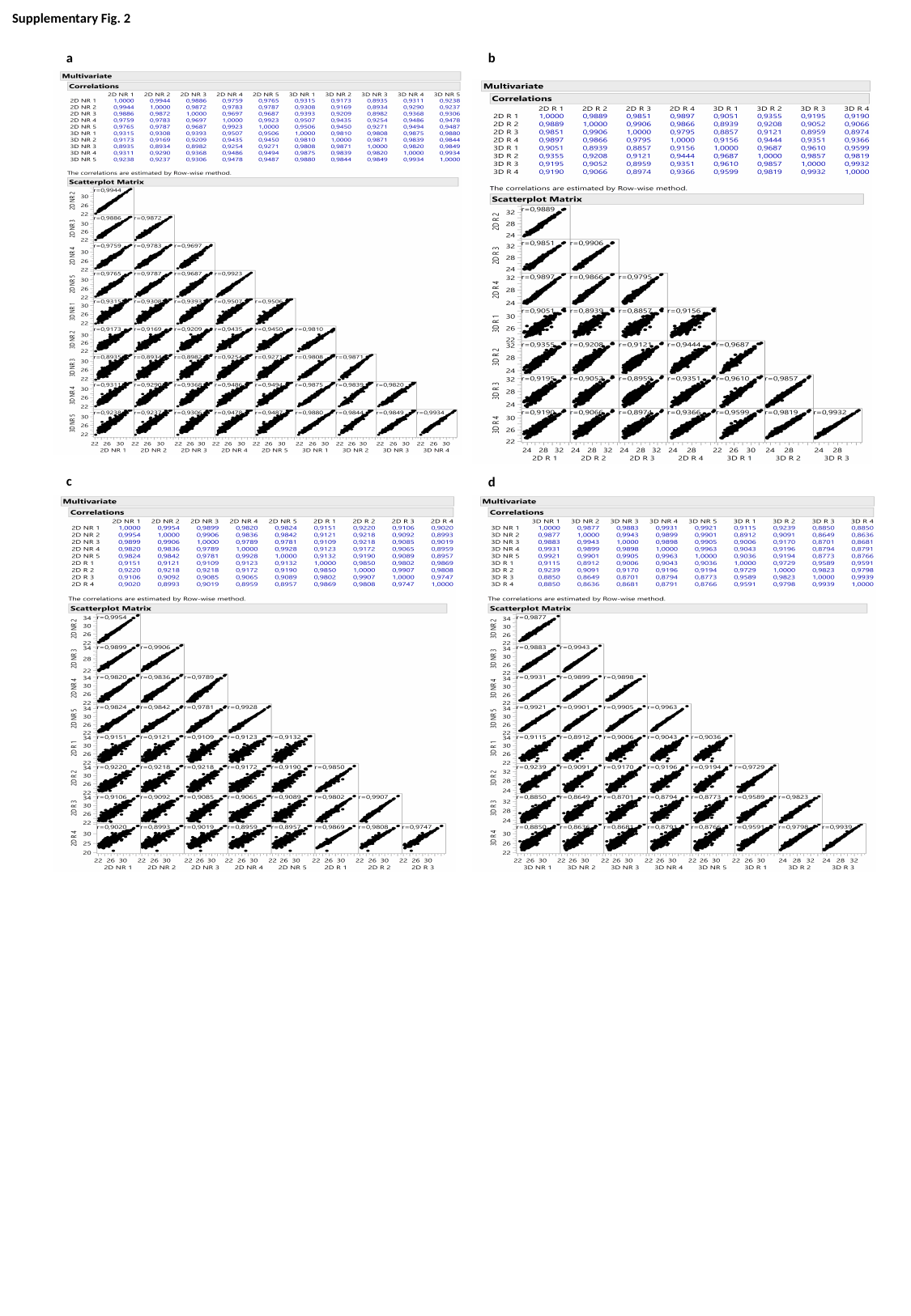

#### Supplementary Fig. 2
a
b
c
d

### Slide 4
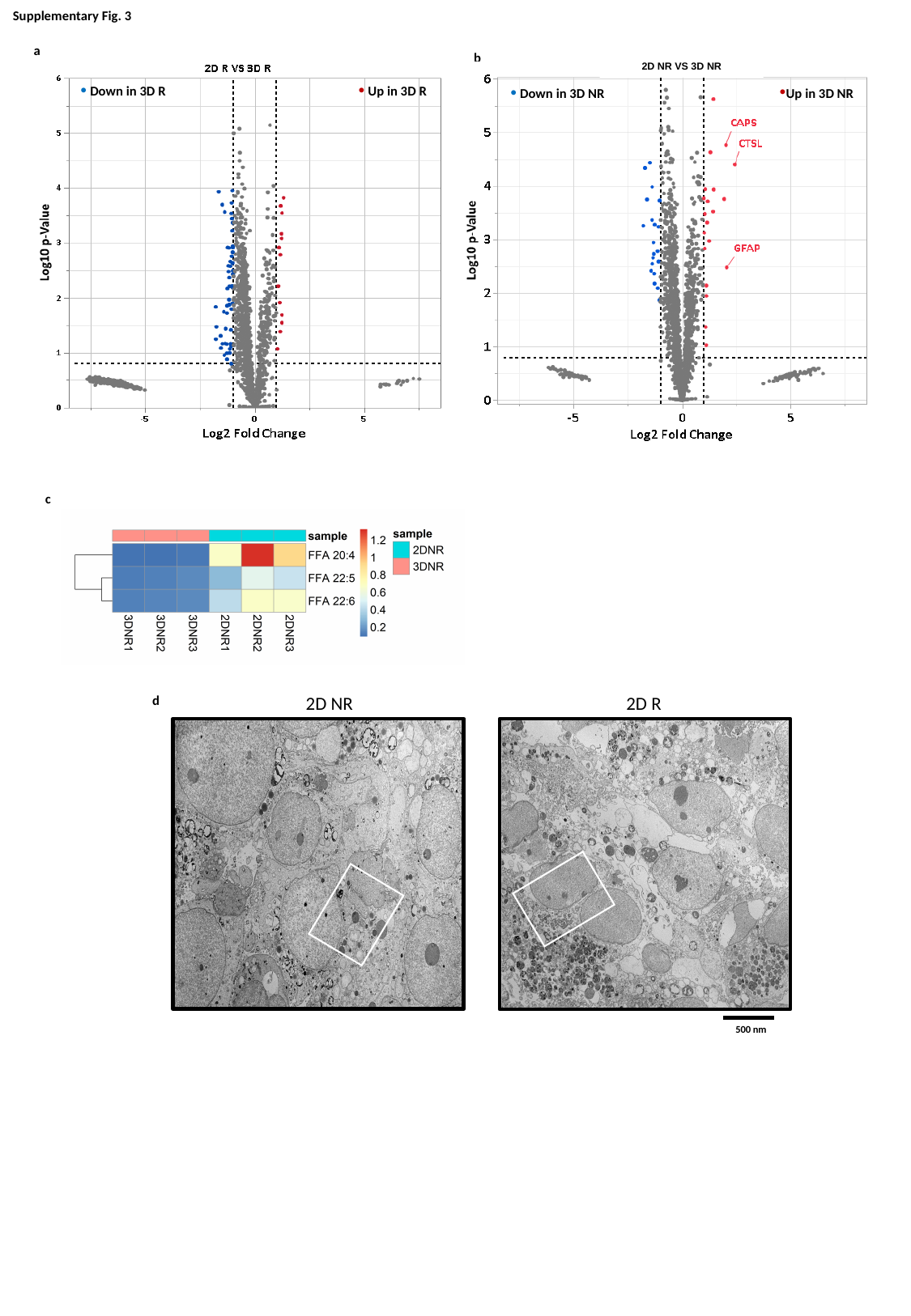

Supplementary Fig. 3
a
 Down in 3D R
 Up in 3D R
b
Up in 3D NR
 Down in 3D NR
2D NR VS 3D NR
c
d
2D NR
2D R
500 nm

### Slide 5
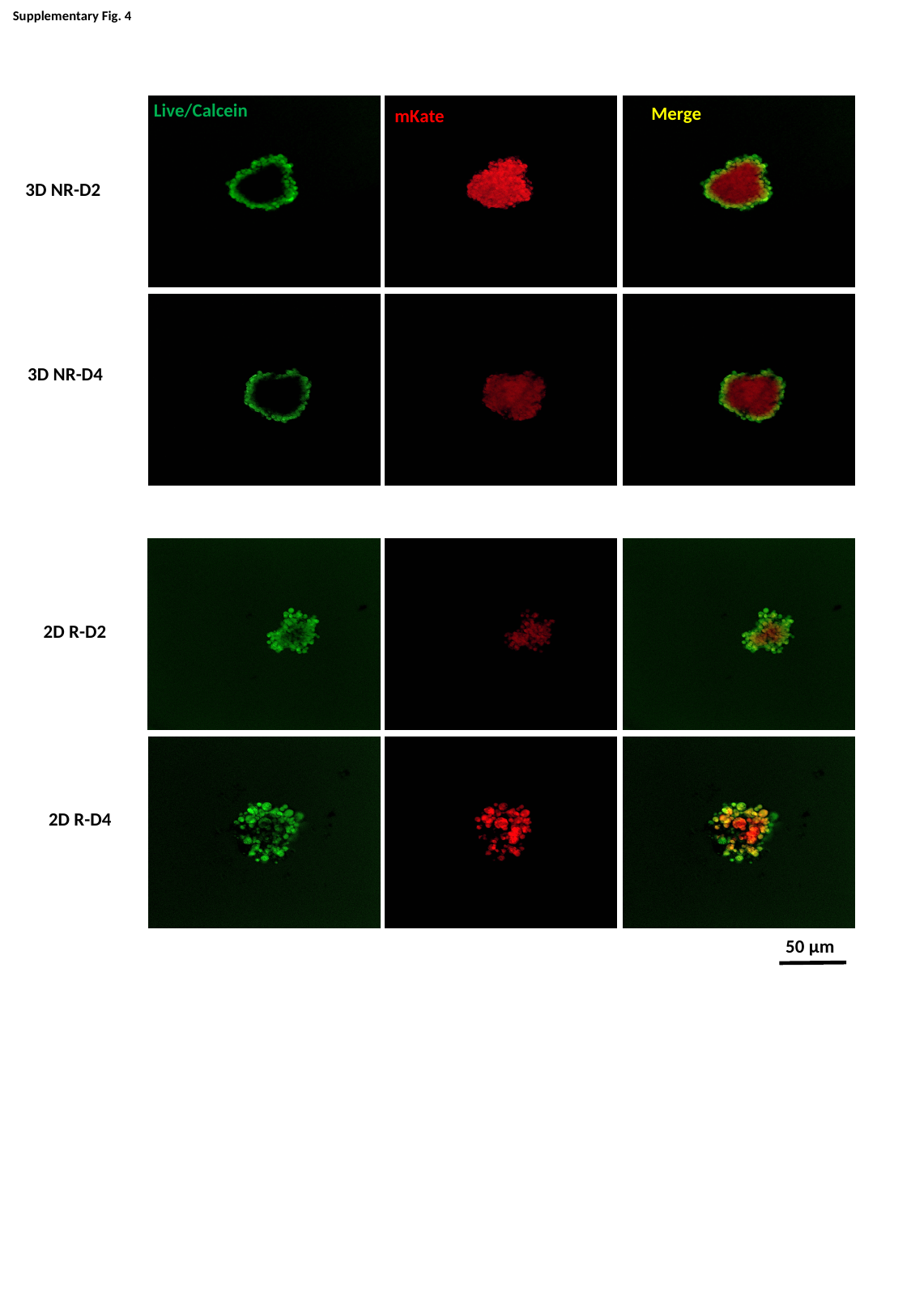

Supplementary Fig. 4
Live/Calcein
Merge
mKate
3D NR-D2
3D NR-D4
2D R-D2
2D R-D4
50 µm

### Slide 6
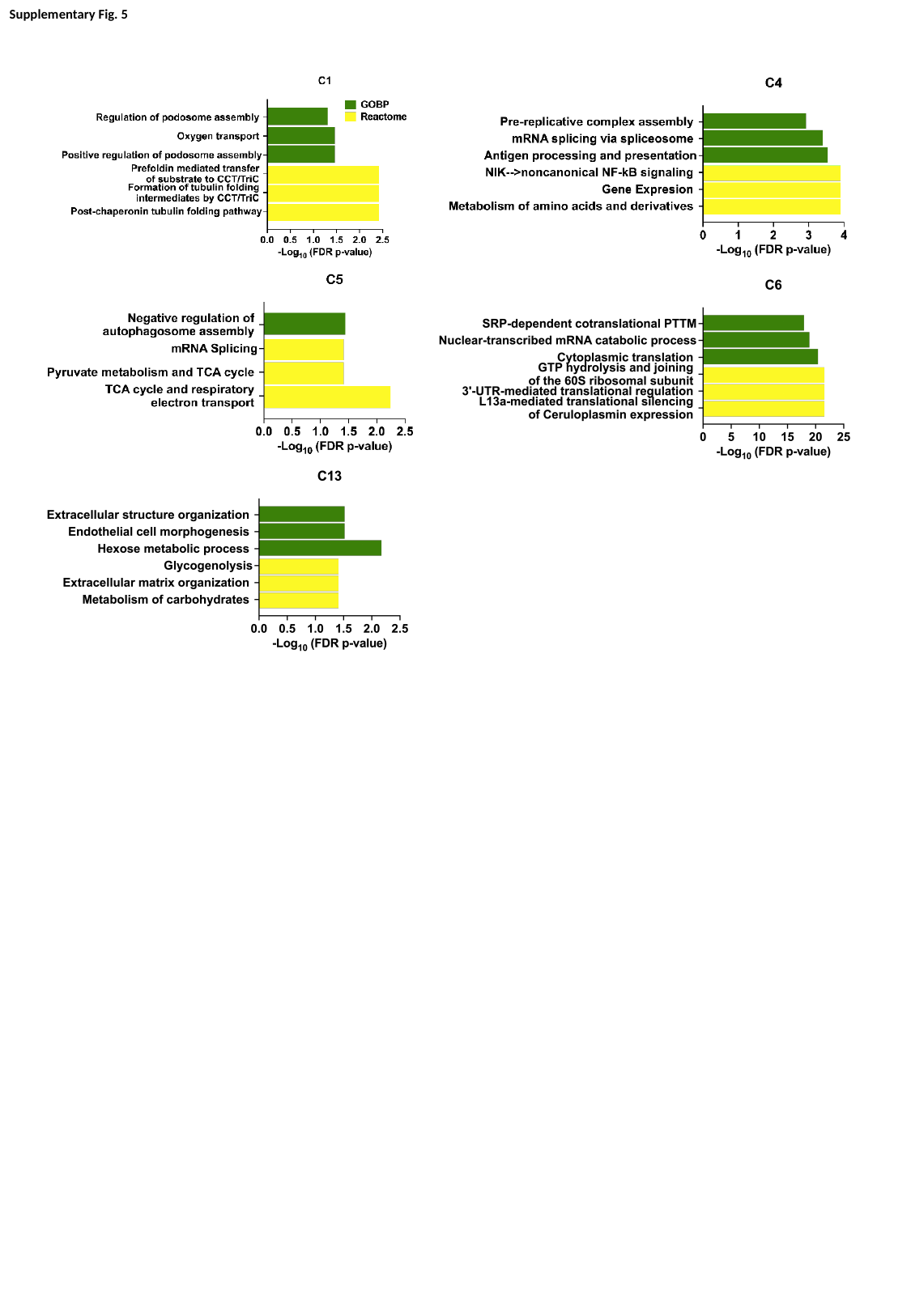

Supplementary Fig. 5

### Slide 7
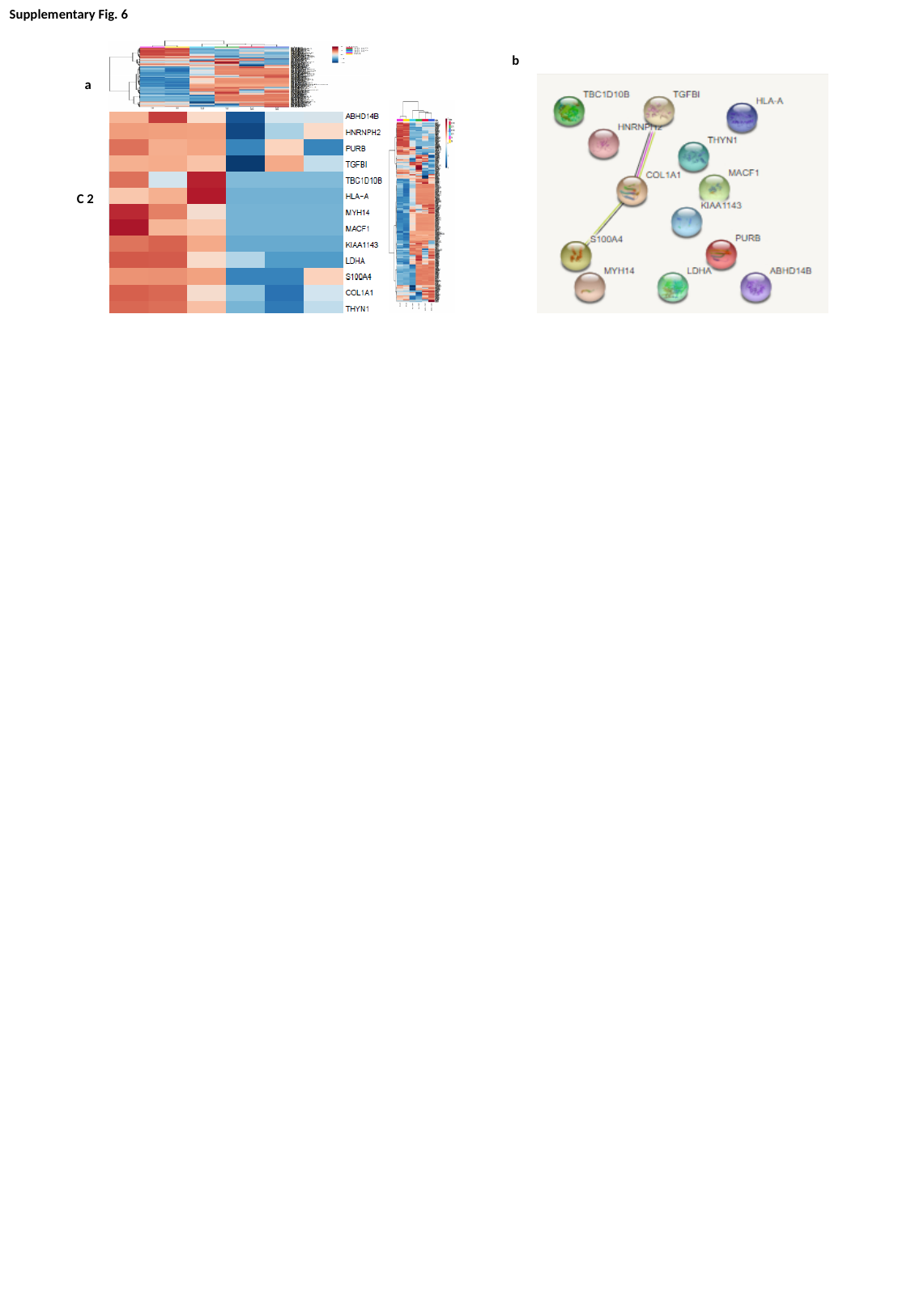

Supplementary Fig. 6
C 2
a
b

### Slide 8
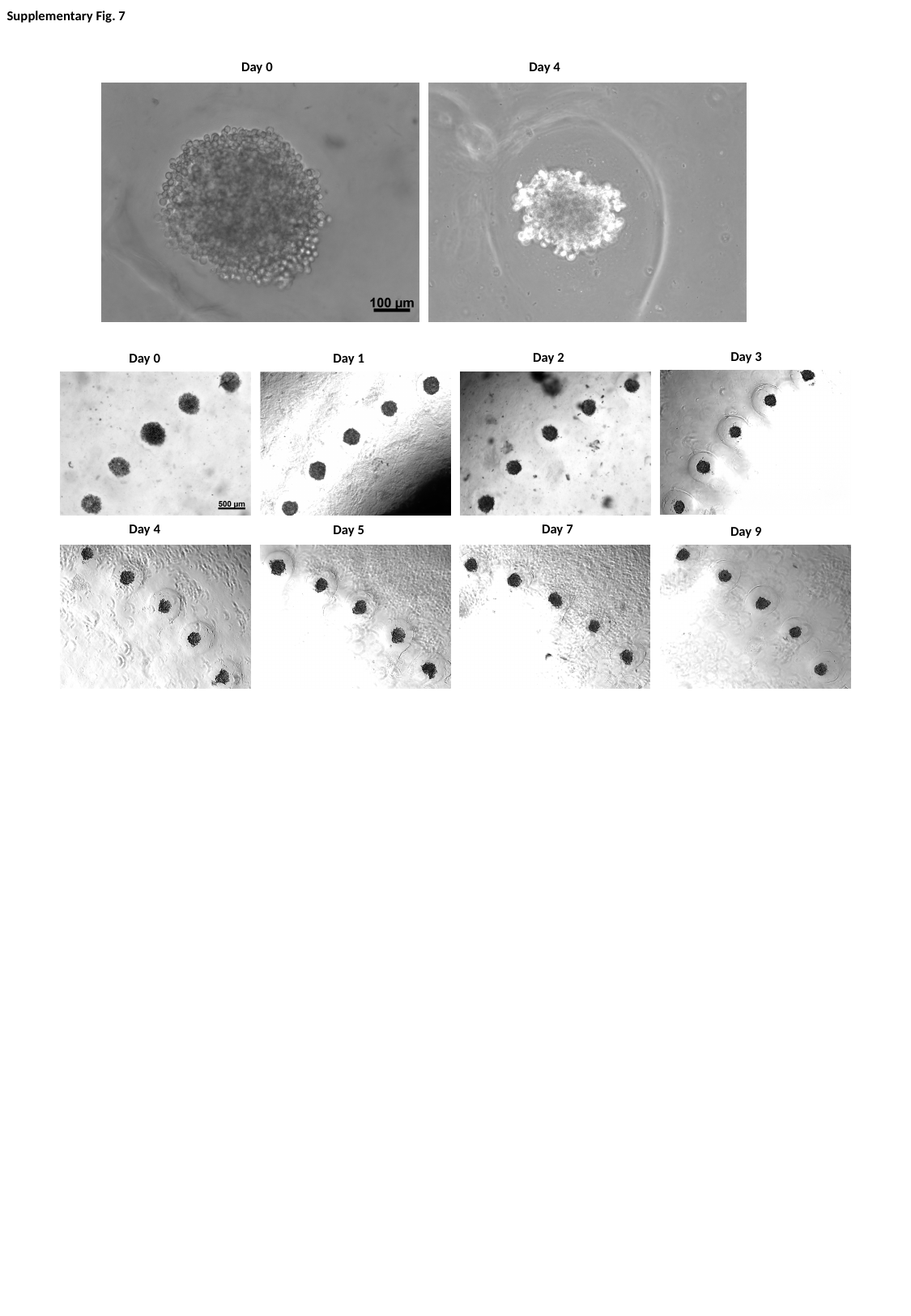

Supplementary Fig. 7
Day 0
Day 4
Day 3
Day 2
Day 0
Day 1
Day 7
Day 4
Day 5
Day 9

### Slide 9
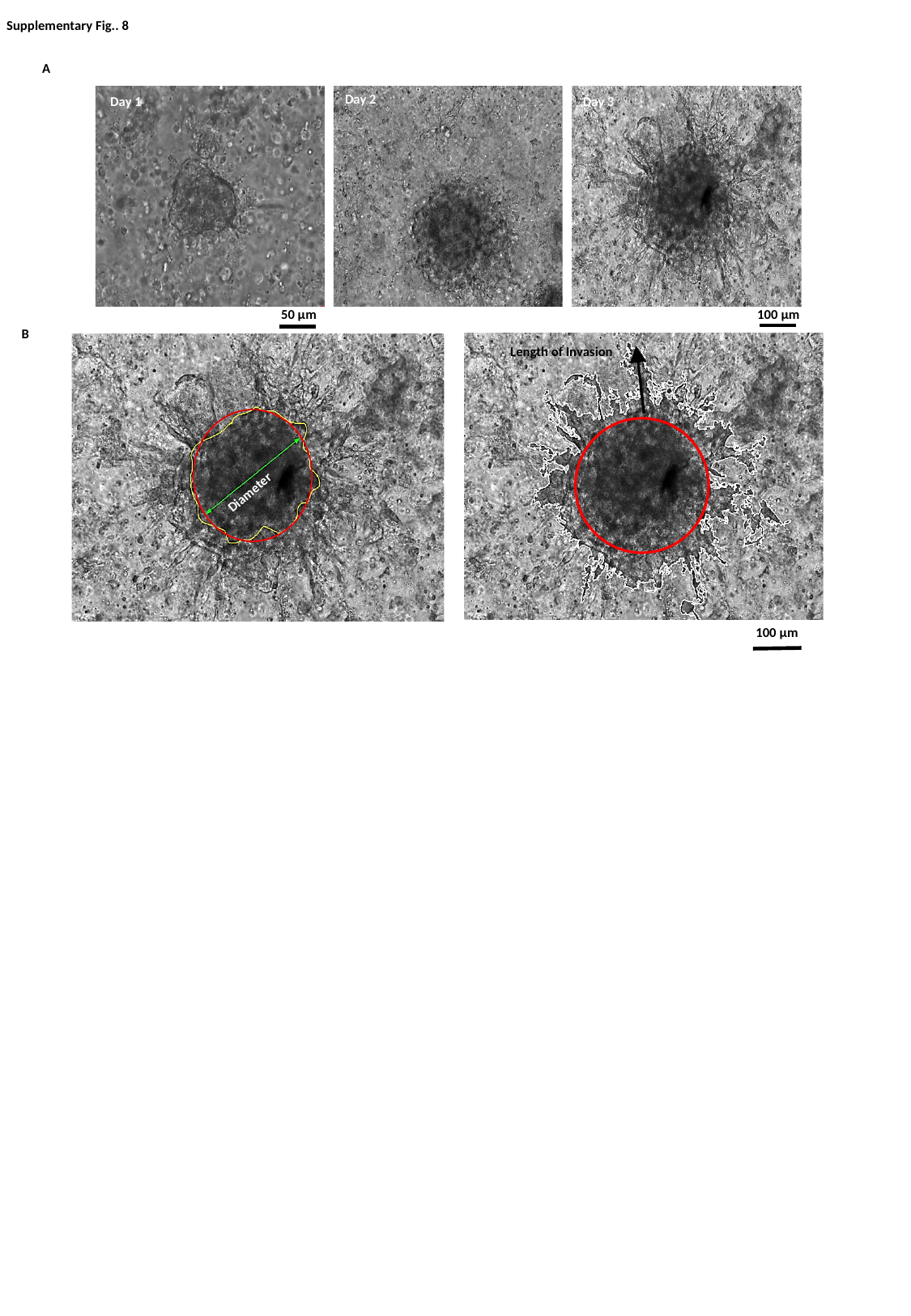

Supplementary Fig.. 8
A
Day 2
Day 1
Day 3
B
Length of Invasion
Diameter
100 μm
50 μm
100 μm

### Slide 10
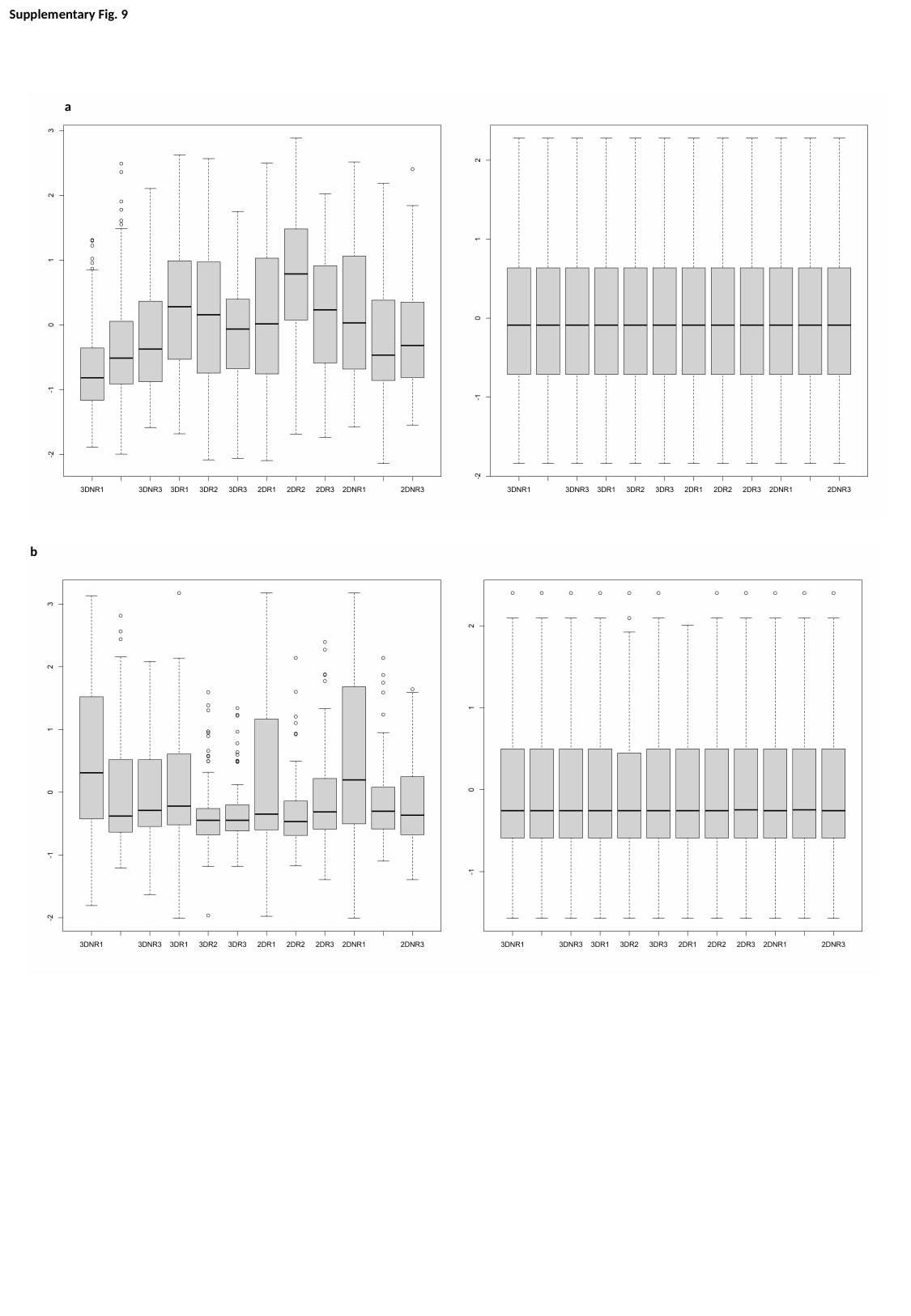

Supplementary Fig. 9
a
b
